# A continuum reaction-diffusion model for spread of gene silencing in chromosomal inactivation

**DOI:** 10.1101/2025.06.02.657345

**Authors:** Shibashis Paul, Sha Sun, Karmella Haynes, Tian Hong

## Abstract

Regulation of gene silencing in large regions of chromosomes is crucial for development and disease progression, and there has been an increasing interest in using it for new therapeutics. One example of massive gene silencing is X chromosome inactivation (XCI), a process essential for dosage compensation of X-linked genes. During XCI, most genes in the X chromosome are inactivated following the transcription of XIST, an X-linked long noncoding RNA. Recent experiments with transgenes showed that the spread of gene silencing can be induced by XIST transcription in *cis*, but the spread is restricted in space. The mechanism of controlling the spread remains unclear. In this work, we develop a continuum reaction-diffusion model that elucidates chromosomal inactivation through a bistable system governed by a regulatory network for XIST-mediated gene silencing. We find that the spread of XIST can be tuned by known negative feedback loops regulating its synthesis and degradation, and that the spread of gene silencing is controlled by a wave-pinning mechanism in which both global regulation of silencing complex and local variations of histone modifications can play crucial roles. In addition, we integrate the discrete three-dimensional arrangement of the X chromosome and autosomes into this continuous model. We use a 3D chromosome structure inferred from experimental data and our modeling framework to show the spatiotemporal regulation for spread of gene silencing. Our method enables the investigation for the inactivation dynamics of large regions of chromosomes with varying degrees of the spread of gene silencing. Our model provides mechanistic insights that quantitatively relate gene regulatory networks to tunability and stability of chromosomal inactivation.

**Author Summary:** Precise control of gene expression is a fundamental process in biology and turning off large parts of chromosomes is both common in many species and important for development and diseases. A well-known example of chromosomal scale gene silencing is X chromosome inactivation (XCI), which helps balancing gene activity between sexes. XCI is governed by the production of an RNA called XIST, leading to most genes on one X chromosome being turned off. Experiments have shown that XIST can trigger nearby genes to turn off in natural and engineered contexts, but this effect only spreads to specific regions. The exact mechanism for this limit is not very well understood. In this study, the authors created a mathematical model to explain how XIST spreads and controls gene activity. They found that feedback systems involving XIST regulations and chromatin modifications synergize and determine how far the inactivation spreads, through a process similar to a wave that gets “pinned” in place. They also included the 3D shape of chromosomes in their model to better understand how gene silencing happens over time and space. This model helps to quantitatively describe chromosome inactivation with gene regulatory networks.

## Introduction

Gene silencing in large chromosomal regions is essential for the development of mammals. Particularly, X-chromosome inactivation (XCI) is used to achieve dosage compensation in female cells to balance the expression of X-linked genes of the two sexes (Payer and Lee, 2008). Before mammalian XCI, one of the two X chromosomes in each cell of the early female embryo is randomly chosen for inactivation. During XCI, the expressions of X-linked genes in the X chromosome selected to be inactivated (Xi) are repressed in a progressive manner upon the transcription of XIST, an X-linked, long-noncoding RNA (lncRNA) (Heard et al., 1997; Kay et al., 1993). XIST molecules are tethered to multiple chromosomal locations, where they recruit polycomb repressive complexes (PRCs) for gene silencing (Sunwoo et al., 2015). The spread of gene silencing was shown to be influenced by the locations of the genes in both one-dimensional (1D) genomic coordinates and three-dimensional (3D) genome structures. The completion of XCI results in the stable repression of most, but not all, genes in the X chromosome (Balaton and Brown, 2016). The number of X-linked genes that remain to be active after the completion of XCI (i.e. the XCI escape gene, or escapees) can vary across tissues and species (Berletch et al., 2015; Tukiainen et al., 2017). The mechanism controlling the extent of gene silencing spread, which in turn determines the escape genes, in XCI remains elusive.

To understand the XIST-mediated gene silencing, multiple groups integrated XIST transgenes into autosomes (Lee and Jaenisch, 1997; Minks et al., 2013; Naciri et al., 2021), and these studies showed the ability of the induced XIST expression in an autosome to silence genes in the adjacency of the XIST integration/transcription site. However, the spread of the gene silencing seems to be limited compared to that in XCI (Minks *et al*., 2013). Similar to the question of determinants of escape genes, factors contributing to the gene silencing spread upon transgene XIST induction are unclear.

Previous mathematical models based on ordinary differential equations (ODEs) describing gene regulatory networks have provided useful insights into the mechanisms of the decision-making process of XCI for robustly selecting one X chromosome in each cell (Li et al., 2016; Mutzel et al., 2019). In addition, agent-based modeling has been used to show the dynamic distribution of XIST molecules over 3D structure of the X chromosome (Lappala et al., 2021). However, these modeling frameworks did not capture the connection between the dynamics of XIST distribution and that of gene silencing. The latter is controlled by complex regulatory networks containing feedback loops in a spatially dependent manner. These feedback loops include the transcriptional and post-transcriptional self-regulations of XIST (Jachowicz et al., 2022; Rodermund et al., 2021) as well as epigenetic modifications that confer gene activity switches and memory (Dodd et al., 2007). The lack of quantitative description of these interconnected regulatory elements has been limiting our understanding of XIST-mediated gene silencing in both natural and engineered systems.

In this work, we used a continuum reaction-diffusion framework to model XIST dynamics and gene silencing induced by XIST upregulation during XCI. We show that the feedback loops of XIST’s transcription and degradation have significant impacts on the steady state distribution of XIST and the speed to attain the distribution. We describe a wave-pinning mechanism based on chromatin-level feedback between histone modifications and repressive complexes, and the proposed mechanism offers a plausible explanation for controllable spread of gene silencing in the X chromosome. In addition, this generic version of our model gives insights into lncRNA-mediated gene silencing in autosomes. Interestingly, both global regulation of silencing complex and local variations of histone modifications can contribute to wave-pinning. We extended the continuum model to capture 3D chromosome structure and showed the utility of our model in explaining key observations such as escape genes under realistic spatial constraints. Overall, our continuum reaction-diffusion model integrates regulatory components contributing to XCI in a systems approach, and it provides new mechanistic insights into factors controlling the spread of XIST-mediated gene silencing.

## Results

### The roles of feedback loops controlling XIST expression on dynamics of gene silencing

To understand how regulations of XIST can influence the XIST distribution, we first built a spatiotemporal model with a one-dimensional (1D) domain and a reaction-diffusion system based on partial differential equations (PDEs). The domain can be viewed as a chromosome region projected onto a 1D space (we will consider a more realistic spatial setting later in this manuscript). A key difference between this model and previous model is that our model allows the diffusion of molecules in continuous space and time, which in turn permits flexibility of molecular movements and descriptions of complex gene regulatory networks. The latter advantage allows us to use a simple set of differential equations to study the effects of molecular concentrations and interactions on gene silencing. Specifically, we first considered a spatial temporal module of regulations for XIST transcription, degradation and diffusion. We used a dimensionless PDE to describe these regulations

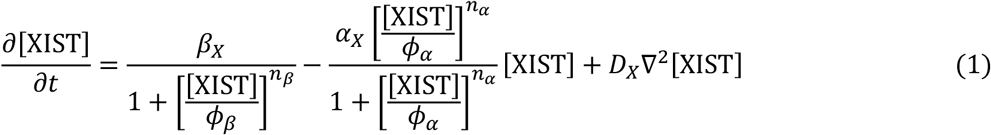

Here, [XIST] represents the concentration of XIST molecules, *βX* is the basal transcription rate of XIST, *αX* is the basal degradation rate constant, *DX* is the diffusion coefficient, *Φβ* is the threshold of the inhibition of XIST’s transcription by itself, *nβ* describes the nonlinearity of this transcription feedback (Jachowicz *et al*., 2022), *Φα* is the threshold of the activation of XIST’s degradation by itself, and *nα* describes the nonlinearity of this degradation feedback (Rodermund *et al*., 2021). Both feedback loops are experimentally observed negative feedback loops (NFLs) in which XIST can limit the level of itself (**Fig 1A**). We selected biologically plausible parameter values to examine the representative dynamical behaviors of the gene regulatory networks (see Supplementary Information). Importantly, the roles of these feedback loops on distributions of XIST and the subsequent gene silencing were unclear due to the lack of rigorous models. To examine NFLs’ function in a quantitative framework, we scanned values of *Φβ* and *Φα* in a grid space with a simple assumption of all other parameters (**Fig 1B and C**). We asked whether the two NFLs can determine how far XIST can reach in the 1D domain with substantial concentration with respect to its level at the transcription site, and how fast XIST distribution can obtain a steady state (**Fig 1B**). Overall, the range of the steady state distribution is negatively correlated with the speed of equilibrium attainment (**Fig 1C**, smaller circles tend to be brighter). However, we found that moderate levels of both feedback gave rise to broad distributions of XIST and relatively rapid attainment of equilibrium (e.g. **Fig 1C** lower right plot. **Figures S1-3**). It should be noted that the broad distribution and rapid equilibrium performance can be achieved simply by increasing the diffusion coefficient of the molecule. However, this way of obtaining the performance would require substantial changes of the physical properties of the molecule (e.g. size). Therefore, our results suggest a powerful approach of tuning the dynamical distribution of XIST by expression regulations alone.

**Figure 1.**
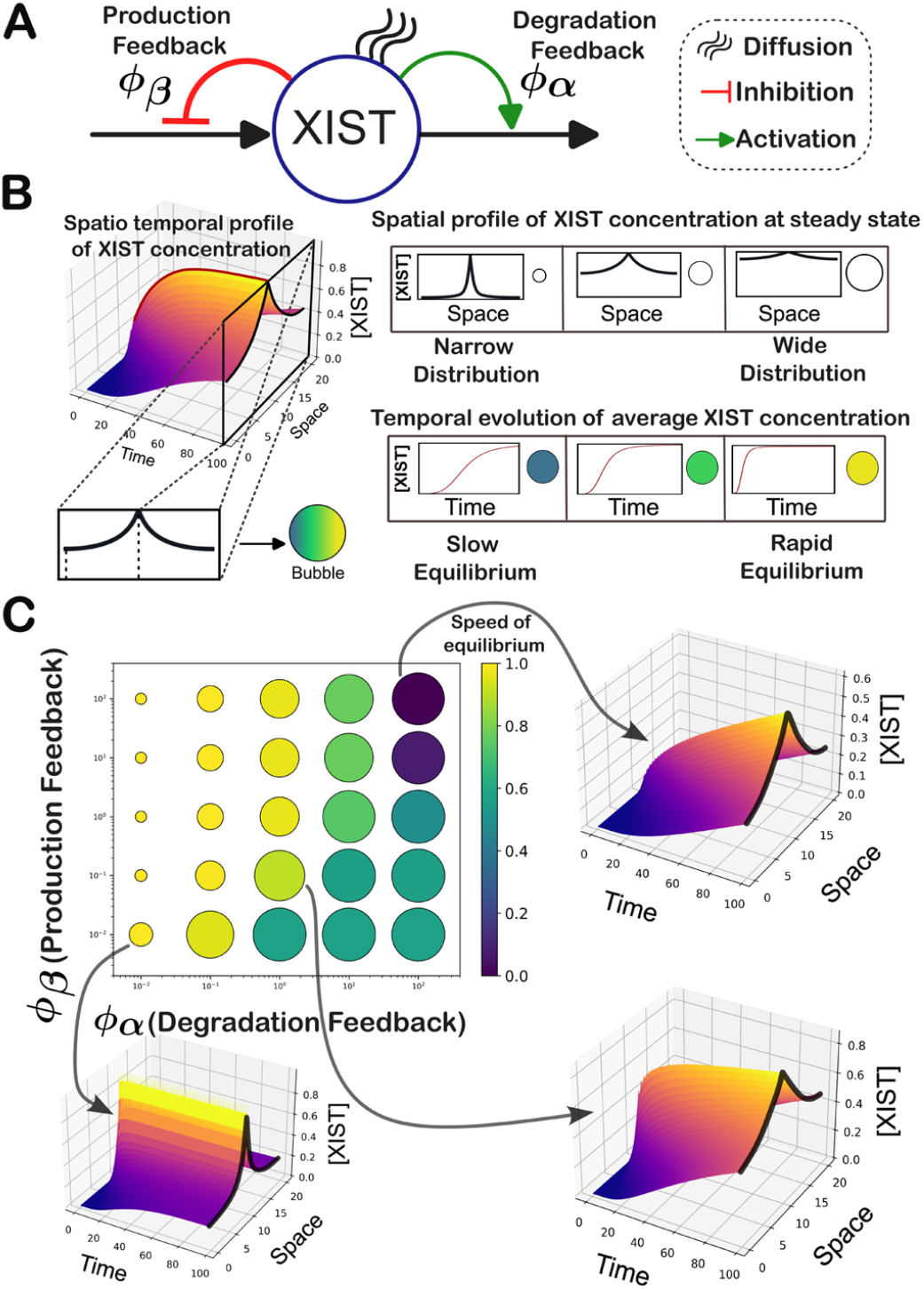
XIST module and its simulation results. A. A regulatory network for controlling XIST levels. This network was used to construct a PDE model to examine the time dependent spatial distribution of XIST. Two feedback loops at degradation and transcription levels are controlled by two threshold parameters *Φα* and *Φβ* respectively. **B**. Simulation of the 1D PDE model (example result shown on left), and two metrics assessing the width and the speed of XIST spread (right). **C**. A grid search for the strengths of the feedback loops (small *Φ* corresponds to lower threshold and stronger feedback) and the associated results of XIST spread width and speed. In these simulations, *nβ* = *nα* = 4, *αX* = *βX* = 1, and *DX* = 1.

### Tunability of the spread of gene silencing by *cis-* and *trans*-acting factors

Next, we focus on a module downstream of XIST that is responsible for the spread of gene silencing. Recent experiments suggest that polycomb repressive complexes (PRCs) 1 and 2, both critical for XCI (Dixon-McDougall and Brown, 2021; Masui et al., 2023), are involved in reciprocal regulations with histone post-translational modifications. Specifically, PRC2 is activated by its enzymatic product H3K27me3 (Stafford et al., 2018). Likewise, histone H2A monoubiquitination (H2Aub) binds and stimulates PRC2 (Kalb et al., 2014), while H2Aub is an enzymatic product of PRC1 which in turn can be recruited by PRC2 (Fischle et al., 2003; Min et al., 2003). These experiments support a positive feedback loop between repressive complexes and the gene-silencing permissive post-translational modifications of histones. We use the following PDEs to describe the dynamics of Polycomb repressive complex PRC2 (as a representative factor for chromatin modification and a transcription repressor) in response to XIST activation:

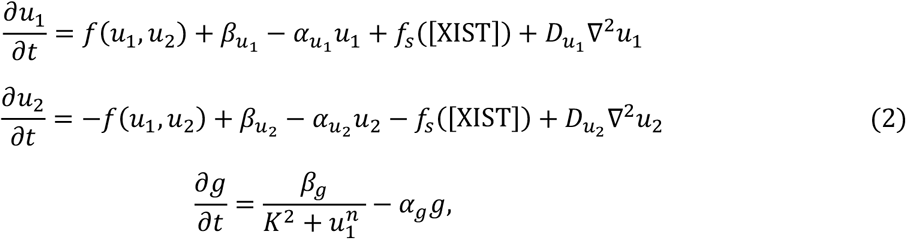

where 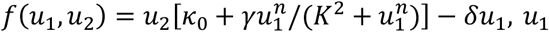 is concentration of the active form of PRC2, *u* 2 is the concentration of the inactive form of PRC2. *∂* represents the activity of any gene in a region of chromosome. *f* (*u* _1_, *u* _2_) represents the interconversion rates between the two forms of PRC2 (**Fig 2A**). The conversion from the inactive form to the active form is assumed to be influenced by the active PRC2 via a positive feedback loop, which is supported by the confirmational change of the complex induced by H3K27me (Stafford *et al*., 2018), a histone modification catalyzed by PRC2. The feedback is characterized by γ, the maximum conversion rate controlled by the feedback, *K* the threshold of self-activation, and *n* the Hill exponent controlling the nonlinearity of the feedback. *κ* _0_ is the basal conversion rate constant from *u* 2 to *u* 1, and *δ* is the conversion rate constant from *u* 1 to *u* 2. 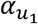 and 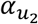 are the disassembly rate of the active and inactive forms of PRC2 respectively. 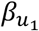 and 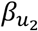 are the assembly rate of the active and inactive forms of PRC2 respectively. 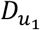 and 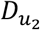 are the diffusion coefficients of the active and inactive forms of PRC2 respectively. The function β *g* / (1 + *u* _1_) describes the rate of gene deactivation triggered by *u* _1_, where *g* is the basal activity. *α g* is the relaxation rate constant of the gene activity. *u s* is the function describing how XIST transiently influences the PRC2 activation (see details below and the rationale for choosing parameter values in the Supplementary Information).

**Figure 2.**
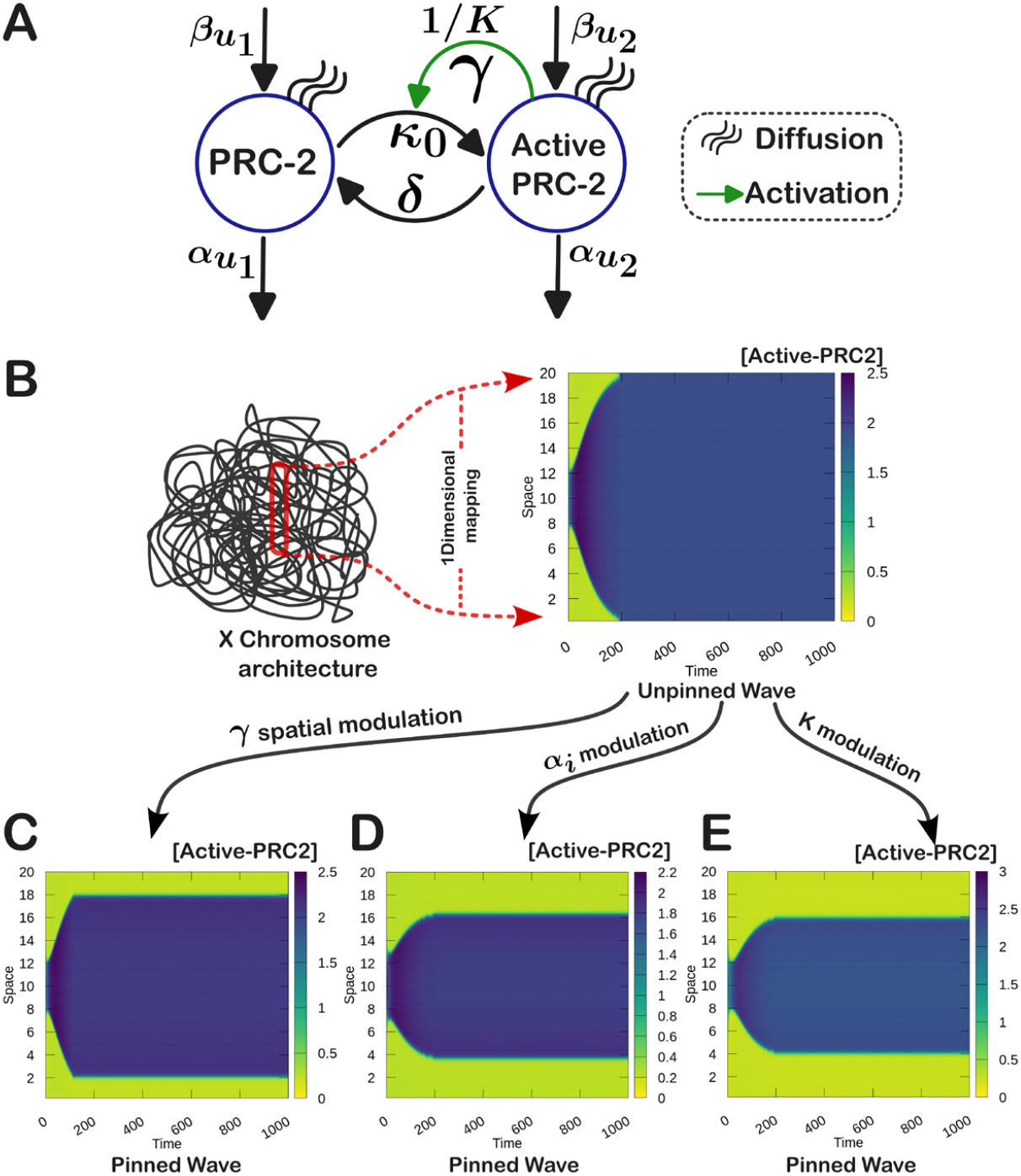
Chromatin module and its simulation results. **A**. A regulatory network for controlling Inactive and Active PRC2 levels. This network was used to construct a PDE model to examine the time dependent spatial distribution of Inactive and Active PRC2 molecules. **B**. Schematic illustration of the mapping procedure of certain chromosomal regions in one spatial dimension. The heatmap represents the time evolution of the Active PRC2 wave leading to homogeneous spatial profile (where, the blue color represents the inactive regions, and the green color represents the active regions of the chromosome). **C**. Attainment of spatial heterogeneity in terms of pinned wave by spatially modulating the more sensitive feedback coefficient γ. **D**. Visualization of the pinned wave achieved through enhancement of degradation coefficient *αX* relative to the unpinned state. **E**. A pinned wave realized by increasing the value of the feedback threshold parameter *K*, which has a lower sensitivity towards the dynamics of XIST molecules. Parameter values for the simulations in this figure are listed in **Table S1**.

To better understand our model in a stepwise fashion, we modularized our model by separately considering the dynamics of PRC2 complex and XIST. We first focus on a *chromatin module* describing PRC2 and gene activities (Eq 2) which does not include XIST explicitly (**Fig 2A**), and *fs* in Eq 2 is assumed to be a transient, imposed stimulation at the beginning of the simulation (e.g. 1 < *∂* < 4) that only occurs in the middle of the spatial domain (see Supplementary Information for details). The dynamics of XIST will be considered more carefully in a later section when we integrate two modules (Eq 1 and Eq 2). With the chromatin module, we simulated the model at a 1D chromosome domain (**Fig 1B**), and we introduced a local perturbation in the middle of the domain (presumably triggered by tethering of XIST to chromatin) where the active PRC2 complex level was elevated while the rest of the region has low a concentration of active PRC2 complex (i.e. a high level of gene expression). As we expected, the positive feedback loop resulted in a trigger wave (also called a traveling wave) of gene silencing (**Fig 2B**), which stems from local bistability of the chromatin states (**Figure S4**). This trigger wave gradually expands the active PRC2-high state to the entire region. Trigger wave driven by positive feedback loop was widely studied in other biological systems (Gelens et al., 2014), and our model suggests a new utility of this mechanism in chromosomal inactivation. Nevertheless, we found that the gene silenced state covered the entire region at steady state even when we simulated a larger domain (**Figure S5**). While this shows the robustness of the trigger wave, the result is inconsistent with the observation that chromosomal inactivation only occurs in one part of the X chromosome during development (Tukiainen *et al*., 2017), and in an even more restricted region when transgene XIST is activated in an autosome (Minks *et al*., 2013; Naciri *et al*., 2021). We therefore asked what can cause the restriction of the spread of gene silencing. Interestingly, reducing the strength of the positive feedback loop in flanking regions of the chromosome resulted in an abrupt termination of the trigger wave (**Fig 2C**). This is consistent with previous reports on sequence and epigenetics determinants for XIST-mediated gene silencing (Loda et al., 2017; Tang et al., 2010). Nonetheless, these local features cannot fully explain the variable extents of the gene silencing spread during XCI and activation of transgene XIST (Tukiainen *et al*., 2017). We therefore tested a global mechanism for wave pinning. By lowering the production and degradation rate constants of PRC2 (α and β in Eq 2), we found that spread of gene silencing can be restricted by a conserved amount of PRC2, which gave rise to a fraction of silenced chromosomal region at the steady state (**Fig 2D**). This global conservation mechanism was similar to the one used for explaining robust polarized distribution of molecules at cell membrane (Mori et al., 2008), and recent theoretical work showed that conservation of the total cellular concentrations of histone modification enzymes is necessary for maintaining epigenetic memory during cell divisions (Owen et al., 2023). We found that the strength of the positive feedback loop is important for the wave pinning (**Fig 2E**), and for the position where the wave is pinned (**Figure S6**). Taken together, our results provide a new mechanistic explanation, i.e. a wave-pinning model, for the spread of gene silencing at the chromosomal level and we identified two distinct factors contributing to the spatial restrictions of the wave. We expect that biological processes such as XCI use a combination of local and global factors for the precise control of gene silencing waves.

### Spread of gene silencing from multiple initiation sites

Previous studies have shown that XIST binds to multiple sites on the X chromosome, and these sites are the places where gene silencing spreads start (Simon et al., 2013). To study potential interactions of multiple silencing waves, we tested the wave dynamics initiated from two sites presumably bound by XIST at the beginning of XCI. Note that here we are still not considering the dynamics of XIST explicitly in this chromatin module. Instead, we introduced two transient stimulations in the simulated spatial domain described in Eq 2. We focused on a wave pinning mechanism based on global control of polycomb repressive complexes because the characteristics of wave pinning driven by local parametric changes are predictable intuitively. Unsurprisingly, with two local stimulations for gene silencing, we observed two silencing waves each starting to spread two both directions in the chromosome region at the beginning of the simulation. However, the two waves became asymmetrical and repelled the propagation of each other (**Fig 3A** left heatmap). To examine the mechanism underlying this phenomenon, we quantified the levels of both active and inactive PRC2 at an early stage of the simulation (**Fig 3A** heatmaps and line plots). When the active PRC2 levels were still symmetrical with respect to the location of the stimulations (**Fig 3A** orange dashed vertical lines), the inactive PRC2 levels started to become asymmetrical: the region between the two stimulations had a lower concentration of inactive PRC2 compared to the exterior regions due to the depletion of inactive PRC2 induced by two waves, as opposed to one wave in each exterior region (**Fig 3A**, green curves).

**Figure 3.**
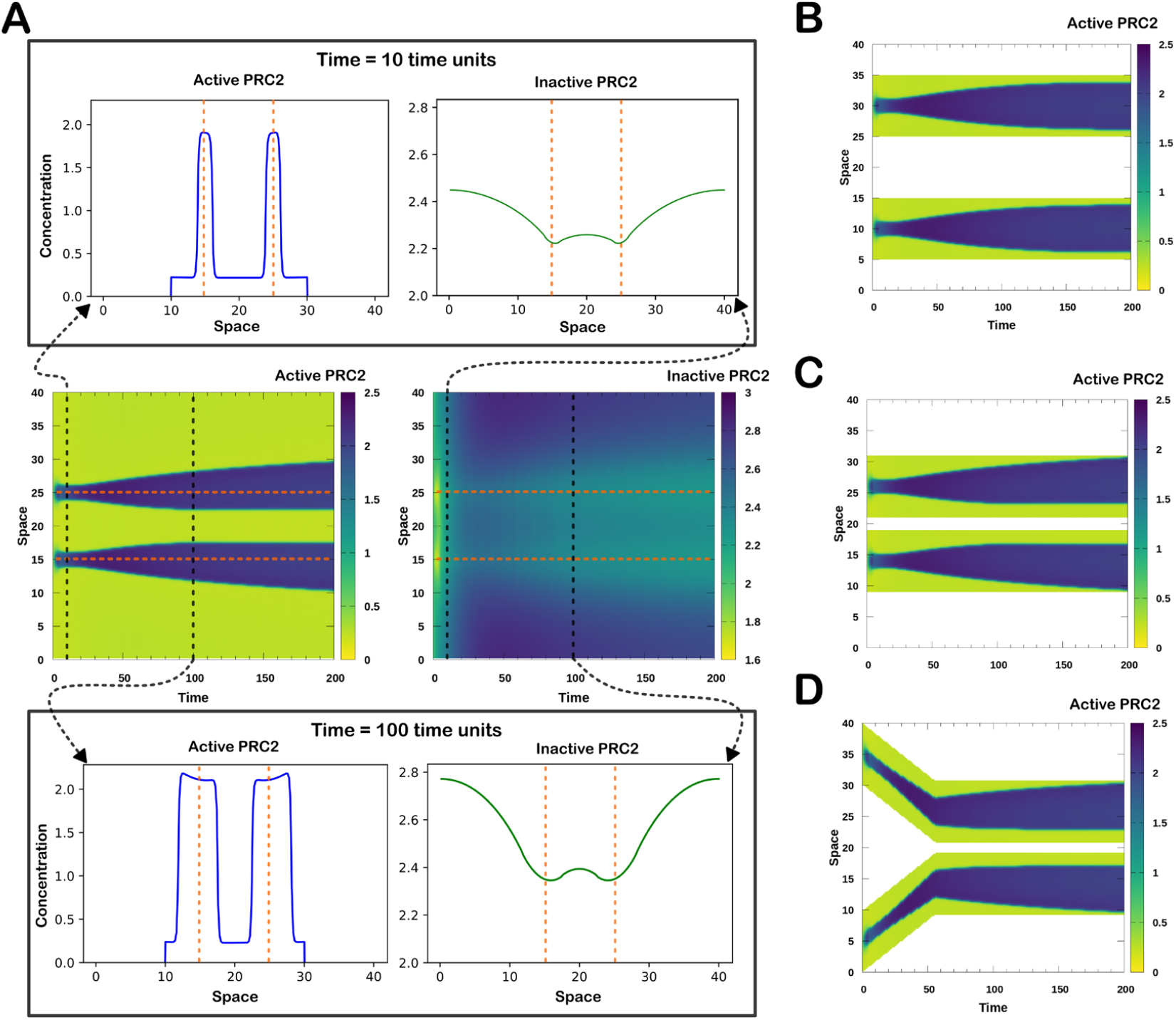
Interaction among the silencing waves and its analysis. A. The heatmaps represent spatiotemporal profile of both active and inactive PRC2 which clearly reveals the repulsive interaction between silencing waves and resulting symmetry breaking. A more detailed analysis of the spatial distribution of both the molecules at two different time points unravels the unequal distribution of inactive PRC2 as the origin of symmetry breaking during the movement of silencing wave fronts. **B-D**. Illustration of the effect of chromosomal movement on spatial profile of active PRC2 complex where the white, blue and green regions represent the void space, silenced and active chromosomal regions respectively. The results indicate that the steady state distribution of the molecules is highly dependent on the final spatial orientation of the chromosome. Parameter values for the simulations in this figure are listed in **Table S2**.

We next asked whether the interactions of silencing waves can influence each other if they were on two separate chromosome regions. Unlike the models presented earlier in this work, the two regions correspond to two chromosomal segments that are not adjacent to each other in genomic coordinates but are relatively close in 3D space. To simulate this in a 1D framework, we created non-chromosomal space that allows diffusion of molecules but cannot be involved in any feedback-related reactions (Eq 2) (white space in **Fig 3B**). When we placed the two segments, each perturbed by a wave-enabling stimulation, with a long distance between them, we found that the two waves did not influence each other’s dynamics (**Fig 3B**). However, when the two segments were close, the two silencing waves repelled each other as we saw with the one-segment model (**Fig 3C**). We next asked whether the movement of chromosomal regions can influence the silencing waves. We simulated a scenario of two chromosomal regions approaching each other, mimicking a chromosome undergoing compaction (**Figure S7**), and we found that the steady state silencing distribution was determined by the steady state, but not transient, positions of chromosomal regions (**Fig 3D**). Furthermore, the repulsion of the two silencing waves was observed for both the no-flux boundary condition, which was used for all simulations presented so far, and the periodic boundary condition (**Figures S8 and S9**). This shows the wide applicability of our conclusions. In summary, our results show that the control of gene silencing via global levels of repressive complexes can give rise to asymmetrical silencing wave propagation, and interactions of multiple waves under both static and dynamic conditions of chromosomal regions’ positions.

### Integration of the XIST and the chromatin modules

We next combined the XIST module (Eq 1) and the chromatin module (Eq 2) to build an integrated system for studying XCI-mediated gene silencing (**Fig 4A**). To bridge the gap between dynamics of XIST concentration and the concentration of the repressor complex, we used an intermediate variable to represent the status of the tethered XIST-protein complex (Pandya-Jones et al., 2020). Higher concentration of XIST triggers the activation of the tethering sites in a probabilistic manner (see Supplementary Information) (**Fig 4B** left and middle panels). The locations of the tethering sites are predefined in the simulated space. The activation of the tethering sites first leads to the activation of the PRC2 complex in the adjacency of the sites, which in turn initiates multiple trigger waves (**Fig 4B** right panel). At the steady state, most of the simulated chromosomal region was silenced, but due to the pinning of the waves, some regions remained to be active, reflecting the stable expression of the genes that have escaped from the chromosomal inactivation. In this model, we assumed spatially homogeneous feedback strengths to examine the effect of global regulations on the silencing wave. Interestingly, some asymmetrical wave patterns were produced due to the random distribution of the tethering sites and the resultant asymmetric interactions among waves that we explained in the previous section (**Fig 3** and **Fig 4**). Overall, the combined model suggests that the XIST-mediated gene silencing can be captured by a dynamical system with multiple traveling waves, and such waves are initiated by both high local concentration of XIST and the existence of local protein assemblies at selected sites of the chromosome.

**Figure 4.**
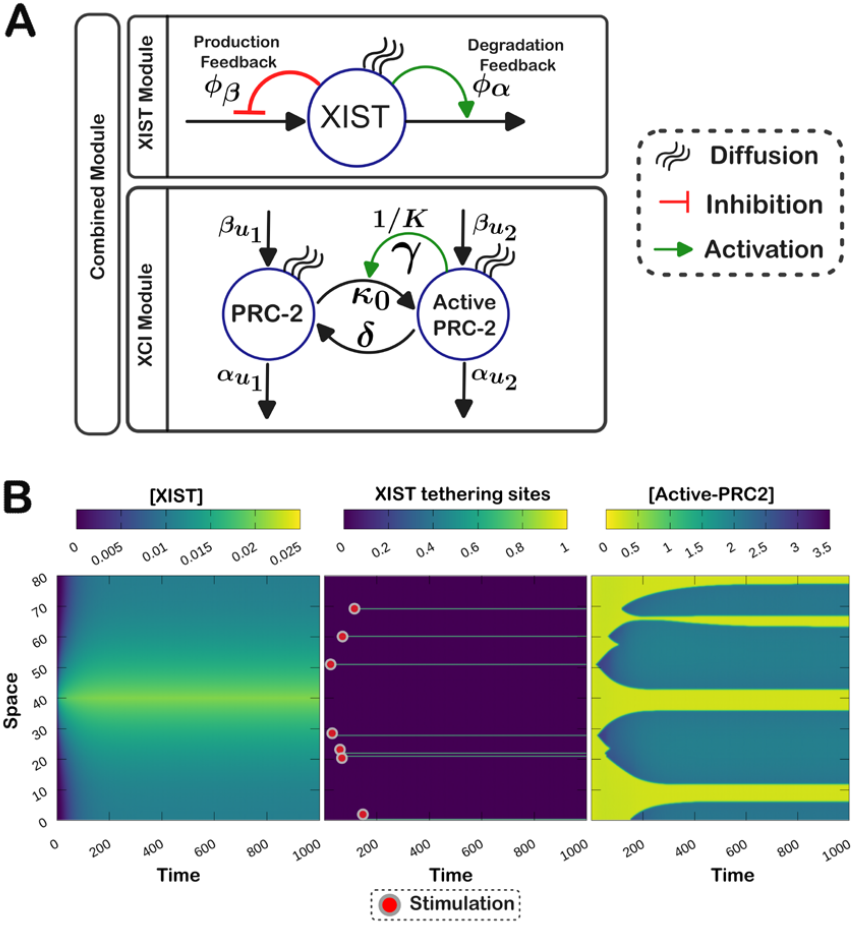
Integrated module and the simulation results. A. Portrayal of the combined biochemical network governing the underlying spatiotemporal dynamics of XIST, active PRC2 and inactive PRC2 molecules. The XIST module and X chromosome inactivation module is bridged by the tethering module that incorporates the effect of XIST spatiotemporal profile to the dynamical landscape of X chromosome inactivation. **B**. The simulation results presented in this sub panel depict the spatiotemporal profile of XIST molecules (left), the tethering sites of the XIST molecules leading to stimulations (center) and the time evolution of the chromosomal inactivation profile (right). A detailed description of the parameters used for the simulation is provided in the Supplementary Information. Parameter values for the simulations in this figure are listed in **Table S3**.

### A 3D model for spatiotemporal dynamics of chromosome inactivation

To illustrate the utility of our modeling framework with realistic chromosome structures and experimental data, we built a proof-of-concept 3D model for X chromosome and its inactivation. We first used a previously published 3D model X chromosome inferred from Hi-C sequencing data (Lappala *et al*., 2021). While the X chromosome structure is dynamic during XCI, we chose the inactive X chromosome structure because we showed with the 1D model that the stationary phase of the chromosome structure is the primary spatial factor determining the silencing waves (**Fig 3**). In addition, since the inferred structure represents an averaged configuration across multiple cells and multiple time points at the stationary phase, we introduced a continuous spatial parameter γ that determines strength of the feedback between chromatin and the repressive complex, reflecting the duration or likelihood of a location in 3D space occupied by chromatin (**Fig 5A, Figure S10**). We used the genomic location of XIST as the production site of XIST RNA, and we used our model the two integrated modules simulate XCI with multiple tethering sites. The location of the tethering sites were fitted to experimental data on silenced genes and escape genes (Berletch *et al*., 2015) (**Fig 5B**). We showed that the model gave rise to a reasonable agreement with experimental data (**Fig 5C-E, Figure S11**) in that only a small fraction of X-linked genes remained active at the steady state and those genes, including XIST itself, were primarily located at the exterior region of the X chromosome in 3D. Among 38 experimentally identified escape genes, the model captured 12 of them while maintaining a nearly 80% silencing percentage of the entire chromosome (**Fig 5E**). Furthermore, we showed that, as a result of the continuous consideration of chromosome structure, silencing waves can spread across multiple chromosomal regions due to their close proximity in 3D (see an example in **Fig 5F** and **G**). This is consistent with a previous experiment using transgene XIST that showed the possibility of gene silencing spread over 3D chromosome structure (Naciri *et al*., 2021).

**Figure 5.**
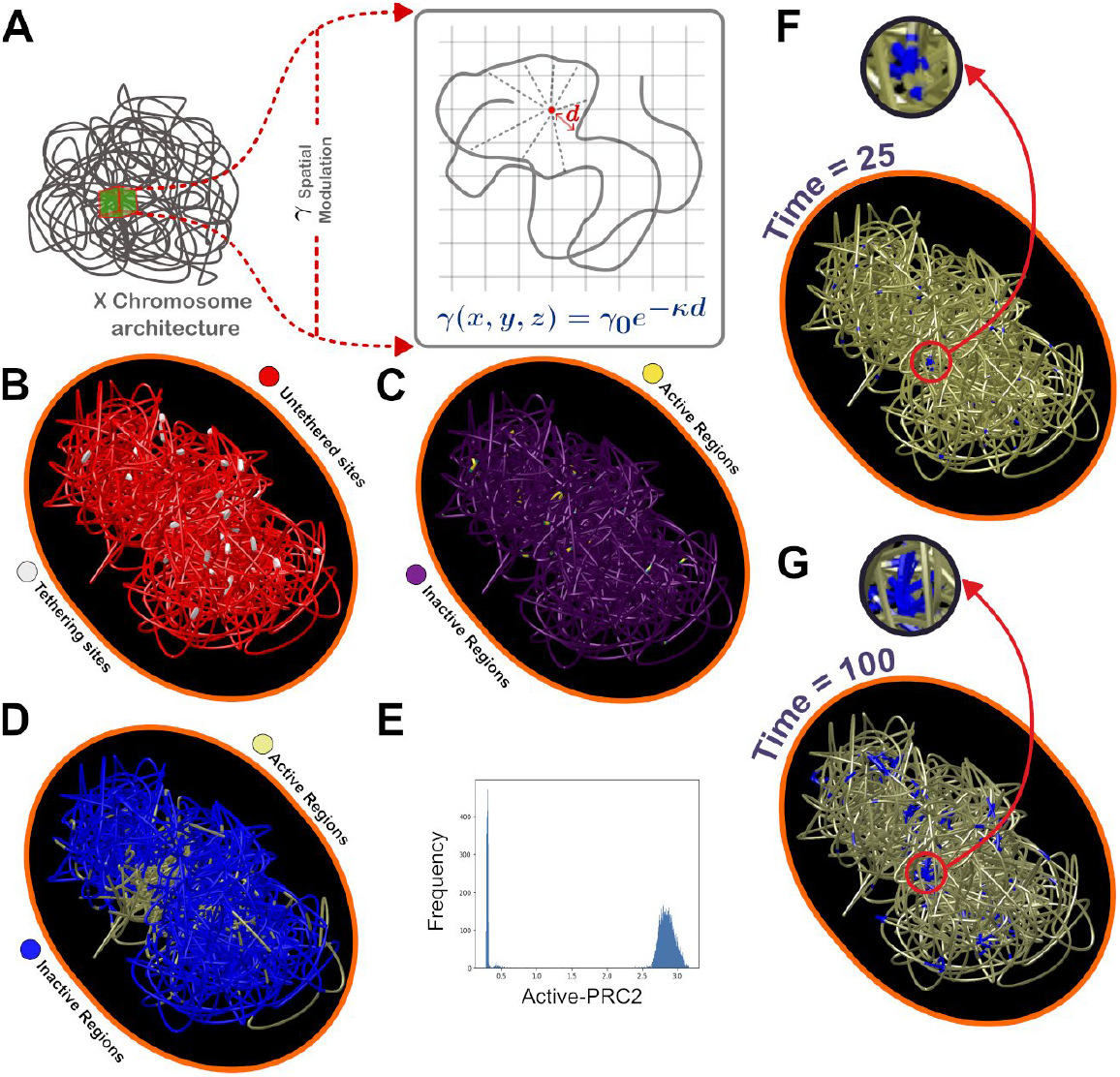
The three-dimensional model and comparison of simulated results with experimental data. A. Schematic representation of implementing spatial dependance of the high sensitivity feedback coefficient γ associated to the process of chromosomal inactivation. **B**. Spatial profile of the XIST tethering sites obtained from 3D simulation marked my white beads. **C**. Spatial profile of the X chromosome inactivation obtained from simulation at time unit 500 which corresponds to approximately 2 hours in physical scale (details about time and length scale estimation are provided in the Supplementary Information file). **D**. Visualization of the experimentally observed active and inactive genes in the X chromosome. **E**. Distribution of chromosomal segments over the levels of active PRC2 in a mouse X chromosome. A total of 20000 evenly spaced points on a simulated chromosome were used. **F-G**. Demonstration of the 3D spatial spreading of the chromosomal inactivation through void space. (The detailed description of the simulation method and utilized parameter values are provided inside the Supplementary Information file, which includes **Table S4**)

## Discussion

X chromosome inactivation (XCI) is a remarkable regulatory program for gene expression in which the transcription of a lncRNA XIST triggers gene silencing for most of the genes in a chromosome. Substantial work has been done to unravel the mechanisms of gene silencing during XCI, but many questions remain open. In particular, it has been unclear how the spread of gene silencing is robustly triggered and regulated in a context dependent manner. In this work, we used a mathematical model based on a continuum reaction-diffusion system to show that feedback loops for XIST transcription and degradation have profound an impact on both the steady state distribution of XIST and the speed to achieve the steady state. We observed that the spatial range of the XIST distribution was increased by transcription-level negative feedback (Jachowicz *et al*., 2022) and decreased by degradation-level negative feedback (Rodermund *et al*., 2021). This suggests that these feedback loops may be used to define the boundary of XIST distribution and its effect on gene silencing, and they may help to limit the XCI to a single chromosome. Previous experimental and theoretical studies without considering spatial distributions showed that negative feedback loop can accelerate the response of gene expression that steady state can be obtained rapidly (Rosenfeld et al., 2002). However, in this work, we showed that when spatial distribution of gene products is considered, transcription-level negative feedback can either speed or slow down response, depending on the choice of other parameters. Interestingly, these functions of negative feedback loops were obtained without changing the diffusion coefficients. This may offer an alternative strategy for cells to tune molecular distributions without significant changes of the physical properties of the molecules through evolution.

The XIST-mediated gene silencing is an example of lncRNA’s substoichiometric action in which a relatively low expression level of a lncRNA can trigger a large effect in gene regulation. It was previously suggested that mechanisms such as molecular condensate formation may explain this type of substoichiometric function (Unfried and Ulitsky, 2022). Our model showed that systems-level positive feedback can amplify the small initial effect induced by lnRNAs and trigger bistable switches (in the case of XCI, an on-to-off switch) in a spatially sequential manner via traveling waves. This is similar to the well-known function of positive feedback for long-range communication without the need of transporting molecules over a long distance (Gelens *et al*., 2014). The local functions of lncRNAs in forming condensates or scaffolding can be combined with downstream positive feedback loops to give rise to large-scale effects, and this type of combination may be common among lncRNA-mediated gene regulation.

Our traveling wave model is based on the assumption of chromatin-level feedback loop between histones and enzymes that catalyze histone modification reactions. This type of feedback was widely used to explain epigenetic memory (Dodd *et al*., 2007). More recently, a polymer-based model showed the importance of both local attraction of epigenetic marks and global regulation of enzymes in establishing epigenetic memory with perturbed 3D chromatin structure (Owen *et al*., 2023). Our models show that imbalanced distributions of polycomb repressive complexes and asymmetrical silencing waves can be a mechanism for local and medium-range attraction and repulsion. The globally driven wave-pinning mechanism is consistent with the finding that global regulation of key enzymes can have significant effects on epigenetic memory.

We used a continuum reaction-diffusion framework for this study, and we the advantages of this framework in describing complex regulatory networks in a spatiotemporal model. This type of network is often difficult to model with agent-based strategies especially when the number of molecules is large. However, a key limitation of the continuum model is that the movement of small numbers of molecules and their dynamic geometries are often not described accurately. Our hybrid approach for modeling 3D chromosome structure in our model is useful for integrating continuous reaction-diffusion and discrete geometry. Nonetheless, we expect that future development of modeling strategies will be necessary to describe the complex system of chromosome-level gene regulation with advantages of both continuum and agent-based methods. Overall, our continuum reaction-diffusion model has provided new insights into mechanisms for gene silencing spread mediated by lncRNAs and it will help to deepen our understanding on gene regulatory networks controlling XCI in both time and space.

## Methods

### Model Development

We have adopted a modular approach to develop our model by dividing the process of chromosomal inactivation into three key steps. First, the spatiotemporal dynamics of XIST molecules, designated as the XIST module. Second, the spatiotemporal dynamics of the inactive and active PRC2 complex, termed the XCI module. Third, the integration of the XIST and XCI (i.e. chromatin) modules, referred to as the Tethering module. Throughout the main and Supplementary Information, we have utilized the above-mentioned terminologies to refer to different modules. The XIST and chromatin modules are represented by three partial differential equations (PDEs) containing production, degradation, reaction, and diffusion terms. Biologically plausible parameter values were chosen to give representative results (Supplementary Information), and scanning of some parameters (feedback strengths) was used to analyze the model behaviors systematically (**Fig 1**). All simulations were performed with the no-flux boundary condition unless otherwise indicated. In a few cases the terms are subject to feedback which is backed by experimental evidence. For integration of the two modules, a bridging Tethering module is represented by a set of conditions depending on the spatiotemporal profile of XIST molecule that invokes the stimulation in XCI module. Importantly, the Tethering module introduces the stochasticity into the model arising out of the limited number of transcribed copies of XIST molecules. In addition to the brief description of each module provided in previous sections, we have presented a detailed description of the same in the Supplementary Information.

### 1D Simulations

After developing and fixing various aspects of the model we proceed further and carry out the phenomenological study of the system by simulating the PDEs in one dimension. We have utilized the explicit forward Euler algorithm implemented in FORTRAN programming language to simulate the PDEs. First, we discretized the space and time and then defined the Laplacian terms in each time step using a central difference formula which is provided in Supplementary Information. A thorough account of numerical simulation and parameters used are provided in the Supplementary Information. We have also implemented the void space, i.e., the region without the presence of chromosome by modulating the feedback coefficient γ, as the parameter has higher sensitivity towards the bistability of the system. Computational details regarding the implementation of void space and spatial movement of the chromosomal region are provided in the Supplementary Information. Finally, we integrated all the modules and carried out the simulation for a wider spatial region and got the steady heterogeneous spatial distribution which is reminiscent to the chromosomal inactivation.

### 3D Simulation

The outcome of the phenomenological studies performed in one spatial dimension required further exploration of the system in three spatial dimensions using a realistic chromosome structure supported by experimental evidence. We have carried out the three-dimensional simulations using a discrete semi-empirical X chromosome structure derived from HiC data. We have also carried out necessary computational maneuvers to simulate our continuum mathematical model over the template of a discrete three-dimensional chromosome structure. The details of the maneuvers are provided in the Supplementary Information. Furthermore, the results from one dimensional study indicated that the spatial profile of the final chromosomal inactivation is explicitly dependent on the final steady geometry of the chromosome. Hence, we have used the stationary structure of inactive X chromosome and performed the numerical simulation in 3D.

### Data Analysis

Subsequently, as a logical extension of our investigation, we have curated the available experimental gene expression data for X chromosome and studied the extent of agreement of the simulated result with the experimental data. Detailed descriptions of conditions to filter the state of a particular gene as on or off are provided in the Supplementary Information.

## Code Availability

Computer code for reproducing simulation results is available at the GitHub repository for this manuscript https://github.com/shibashispaul32/XCI.

## Acknowledgements

This work is supported by grants from National Institutes of Health (R35GM149531 awarded to T.H.) and from National Science Foundation (2243562 awarded to T.H. and 2350447 awarded to K.H. and T.H.). The authors thank Rachel Patton McCord for her critical reading of this manuscript.

