## Supplementary Information for "A continuum reaction-diffusion model for spread of gene silencing in chromosomal inactivation"

[2]Department of Developmental and Cell Biology. University of California, Irvine

[3]Wallace H. Coulter Department of Biomedical Engineering. Emory University

### 1 Introduction

X chromosome inactivation (XCI) is a fascinating biological phenomenon that achieves the equalization of X chromosome-linked gene expression between male and female cells. It operates akin to a biochemical switch, deactivating redundant X chromosomes in female cells. Investigating XCI serves as a crucial avenue for comprehending large-scale gene silencing processes. While conventional particle-based statistical modeling has been instrumental in studying XCI, it encounters challenges in addressing certain biochemical aspects, particularly the intricate feedback mechanisms associated with XCI, due to their computational complexity. As a result, this paper presents a continuum model based on partial differential equations for elucidating the gene silencing process.

### 2 The Model

We have developed a partial differential equation-based reaction diffusion model to study the process of XCI. Using the partial differential equation framework has equipped our study with several advantages, such as implementing reaction kinetics with complex biochemical feedback mechanisms and computational efficiency, to name a few. In contrast, in the initial phase, our study suffered from a striking disadvantage of using a continuum formulation, i.e., incorporating the effect of the intricate three-dimensional architecture of the X chromosome on the process of XCI. Nevertheless, by utilizing the reaction kinetics of X chromosome inactivation, we have managed to include the chromosomal structure into our model, which we will discuss in the later part of this paper. We have developed our model in a modular approach, dividing the process of XCI into two different modules. Once we have standardized each individual module, we combine them to build the complete model. The process of XCI can be divided into two major modules, namely, (1) spatiotemporal dynamics of the key biochemical player of XCI, a long non coding RNA called XIST, and (2) spatiotemporal dynamics of XIST-mediated methylation of the nucleosomes. Once we have settled with both modules through appropriate parameter optimization, we combine them through another module, namely, (3) XIST tethering to the X chromosome. Detailed descriptions of these modules will be provided in the following subsections.

#### 2.1 Spatiotemporal dynamics of XIST molecules (XIST module)

Recent experimental studies reveal that XIST molecules exhibit negative auto-regulation on their own production and positive auto-regulation on their own degradation (Fig.1A). In our modeling framework, we have aimed to capture this feedback mechanism by formulating the following partial differential equation originating from mass action kinetics,

$$\frac{\partial[XIST]}{\partial t} = \frac{\beta_X}{1+\left[\frac{[XIST]}{\phi_\beta}\right]^{n_\beta}} - \frac{\alpha_X[XIST]}{1+\left[\frac{[XIST]}{\phi_\alpha}\right]^{-n_\alpha}} + D_X\nabla^2[XIST] \tag{S.1}$$

Where, the first and second terms constitute two Hill functions and are representing production, and degradation of XIST molecules respectively. The third term constitutes a Laplacian and it is representing the spatial diffusion of the XIST molecules. Furthermore, $\beta_X$ is the basal transcription rate of XIST RNA molecules, $\alpha_X$ is the basal degradation rate of the same; $\phi_\beta$ and $\phi_\alpha$ are the negative and positive feedback coefficients, respectively for XIST production and degradation. $n_\beta$ and $n_\alpha$ describe the corresponding Hill coefficients. We have investigated the effect of feedback mechanisms on the spatiotemporal profile of XIST molecules which we will be discussed in later sections.

#### 2.2 Spatiotemporal dynamics of chromosomal inactivation (Chromatin module)

During X chromosome inactivation (XCI), the long non-coding RNA XIST spreads throughout the inactive X chromosome in cis, eventually tethering to specific regions of the chromosome's architecture. Once tethered, XIST molecules recruit the Polycomb Repressive Complex 2 (PRC2), a reader-writer enzyme, and transition its state from inactive to active. The activated PRC2 enzyme then initiates gene silencing by methylating the nucleosomes.

Recent experimental findings indicate that the rate of transition of PRC2 from an inactive to an active state is positively regulated by neighboring inactive epigenetic markers, specifically of the type H3K27Me3.

Based on these biological processes, we have developed a set of partial differential equations with embedded bi-stability to describe the spatiotemporal dynamics of the active and inactive forms of the PRC2 enzyme.

$$\frac{\partial u_1}{\partial t} = f(u_1, u_2) + \beta_{u_1} - \alpha_{u_1} u_1 + f_s([XIST]) + D_{u_1} \nabla^2 u_1 \tag{S.2}$$

$$\frac{\partial u_2}{\partial t} = -f(u_1, u_2) + \beta_{u_2} - \alpha_{u_2} u_2 - f_s([XIST]) + D_{u_2} \nabla^2 u_2 \tag{S.3}$$

$$f(u_1, u_2) = u_2 \left[ \kappa_0 + \frac{\gamma u_1^n}{K^n + u_1^n} \right] - \delta\, u_1 \tag{S.4}$$

$$u_1 = [\text{Active-PRC2}];\ u_2 = [\text{PRC2}]$$

Where, $u_1$ stands for the active form of PRC2 and $u_2$ the inactive form. As the process of chromosomal inactivation is carried out by active PRC2, hence in the context of our model the higher concentration of $u_1$ represents the inactivation of chromosome and vice versa. $\beta_i$, $\alpha_i$ and $D_i$ portrays respectively the production rate and degradation rate and rate of diffusion of corresponding species. $\kappa_0$ is the rate of switching from the inactive to active form and $\delta$ is the rate of the reverse switching. $\gamma$ and $K$ both modulate the positive feedback imparted by the neighboring epigenetic markers with different degree of sensitivity. $n$ is the Hill coefficient. $f_s$ is the stimulation function which is extensively dependent on the spatiotemporal profile of XIST molecules and probability of tethering. The detailed description for the dependency of $f_s$ on the spatiotemporal profile of XIST molecules will be discussed in the next subsection where we have addres@articleID, author = author, title = title, journaltitle = journaltitle, date = date, OPTtranslator = translator, OPTannotator = annotator, OPTcommentator = commentator, OPTsubtitle = subtitle, OPTtitleaddon = titleaddon, OPTeditor = editor, OPTeditora = editora, OPTeditorb = editorb, OPTeditorc = editorc, OPTjournalsubtitle = journalsubtitle, OPTissuetitle = issuetitle, OPTissuesubtitle = issuesubtitle, OPTlanguage = language, OPToriglanguage = origlanguage, OPTseries = series, OPTvolume = volume, OPTnumber = number, OPTeid = eid, OPTissue = issue, OPTmonth = month, OPTpages = pages, OPTversion = version, OPTnote = note, OPTissn = issn, OPTaddendum = addendum, OPTpubstate = pubstate, OPTdoi = doi, OPTeprint = eprint, OPTeprintclass = eprintclass, OPTeprinttype = eprinttype, OPTurl = url, OPTurldate = urldate, sed the XIST tethering module.

We estimated biologically plausible ranges of parameters for our model. The PRC2 complex primarily consists of Polycomb Group (PcG) proteins, including EZH1/2, EED, and SUZ12 [1]. Since experimental data on the overall production rate of PRC2 are currently unavailable, we have assumed that its synthesis rate is correlated with that of PcG proteins and operates within their biologically permissible synthesis range. Reported turnover rates for PcG proteins typically range between 1 and 1000 molecules per hour per cell considering the somatic cells as well as cancerous cells [2, 3]. Assuming homogeneous distribution and typical mammalian cell volume of $\approx 2\ pL$ [4] leads us to the acceptable range of synthesis rate 0.1-100 $pM/sec$. Similar calculation reveals the range of degradation rate to be 14-140 $pM/sec$ [5]. We used a representative cooperativity level ($n = 4$) for the Hill functions in our models. There are a variety of biological mechanisms that can support ultrasensitive responses, and this choice of parameter reflects the high nonlinearity underlying the common biological phenomenon.[6, 7]

#### 2.3 XIST tethering to the X chromosome (Tethering module)

After standardizing the previous modules, we integrate them with a third module called the XIST tethering module to develop a comprehensive model for X Chromosome Inactivation (XCI). Within this module, we have incorporated spatial tethering sites for XIST molecules. The tethering process at a specific spatial point is governed by two key factors. First, the concentration of XIST molecules, which is derived from the simulation of Eq. S.1. Second, the tethering probability, defined as the ratio of the maximum allowed tethering sites ($n_{\text{thr}}$) to the total number of available sites for tethering in the chromosome structure ($n_{\text{total}}$). Therefore, by combining these conditions, we can express the tethering probability ($P_{\text{thr}}$) at any given spatial co-ordinate $\mathbf{q}$ and time point $t$ as:

$$P_{\text{thr}}(\mathbf{q}, t) = \begin{cases} 1 & \text{if } [XIST](\mathbf{q}, t) > [XIST]_C(t) \text{ and } r < \frac{n_{\text{thr}}}{n_{\text{total}}} \\ 0 & \text{otherwise} \end{cases} \tag{S.5}$$

Where, $r$ is a random variable with uniform distribution, spanned between 0 and 1 and $[XIST]_C(t)$ is the time dependent critical concentration for tethering of XIST molecules to the X chromosome and we have defined it as follows:

$$[XIST]_C(t) = [XIST]_{\min}(t) + \phi_{\text{thr}} \left\{ [XIST]_{\max}(t) - [XIST]_{\min}(t) \right\} \tag{S.6}$$

where, $[XIST]_{\min}$, and $[XIST]_{\max}$ is the minimum and maximum XIST concentration at time $t$, respectively. $\phi_{\text{thr}}$ is the tethering coefficient (as the spatial profile of XIST molecules is time dependent so the critical XIST concentration for tethering should also be time dependent) which controls the susceptibility of the XIST molecules for tethering to the chromosome and this parameter should be fine tuned based upon the spatiotemporal profile of XIST molecules. In summary, the tethering module determines the initial stimulation points for the XCI module by utilizing the simulation results from the XIST module.

### 3 Methods

We conducted our studies in a manner reminiscent of our model development, where we performed numerical simulations of partial differential equations chronologically. Initially, we focused on the XIST module, conducting several numerical simulations to uncover the determining factors influencing the spatiotemporal profile of XIST molecules. Subsequently, we systematically explored the XCI module to incorporate biologically relevant features and identify determining factors affecting the extent of chromosomal inactivation. Thereafter, we incorporate the tethering module into our simulation and conduct a thorough computational investigation of the entire model. Moreover, we have investigated the effect of chromosomal compaction on the final profile of X chromosome inactivation in the context of our model. Up until this point, we have performed the simulation of our model considering only one spatial dimension to unravel salient features of our model in a phenomenological manner. Finally, we have implemented our model in three spatial dimensions including all the features we have discussed thus far and compared the simulation result with experimental results. In this section we will present a detailed description of several aspects of the simulation methods that we have utilized to obtain the results.

#### 3.1 Simulations performed in one dimension

##### 3.1.1 XIST module

We have performed numerical simulation of the XIST module, i.e., partial differential equations described the Eq.S.1. We have used the explicit Euler algorithm for numerical integration of the PDEs and used central difference formula to evaluate the Laplacian for a particular spatial point at each time steps. The central difference formula for one dimensional Laplacian can be written as:

$$\nabla^2 u(x) = \frac{u(x+\Delta x) + u(x-\Delta x) - 2u(x)}{\Delta x^2} \tag{S.7}$$

Here, $u$ is the arbitrary species of concern and $\Delta x$ is the spatial grid size.

We simulated the XIST module across various parameter sets to identify the optimal feedback strength for XIST production and degradation ($\phi_\beta$ and $\phi_\alpha$ respectively), ensuring the most rapid and homogeneous spatial spreading of XIST molecules over time. The results are depicted in the following two figures: A pertinent point to be noted is that for the simulations we have set $\beta = 1$ for X inactive center, i.e., the transcription site of XIST and $\beta = 0$ otherwise. The Figs.S1 and S2 together reveal that there is an optimal value of feedback coefficients ($\phi_\beta$, $\phi_\alpha$) for which the XIST

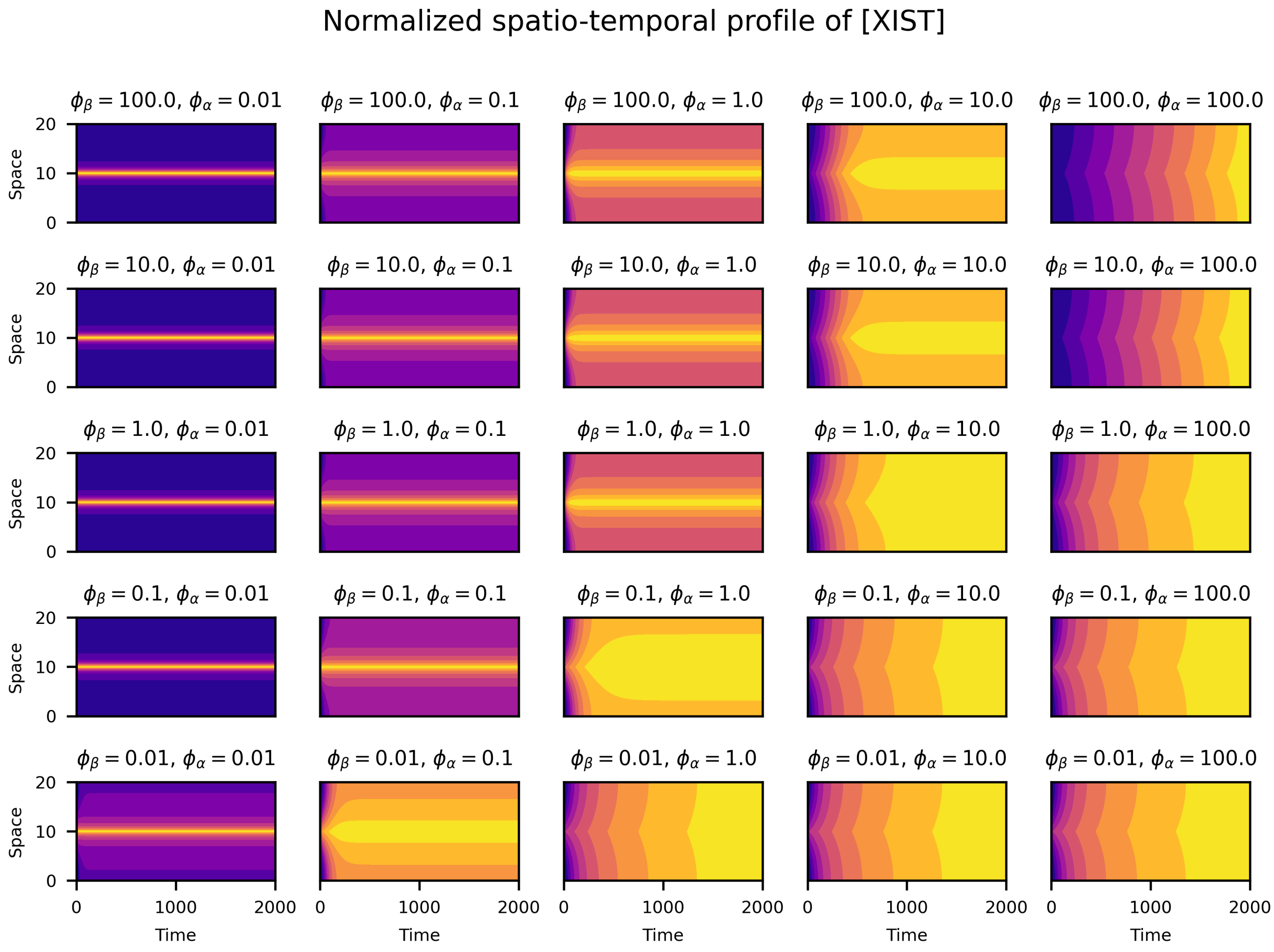

Fig. S1: Spatiotemporal profile of XIST concentration for different values of feedback coefficients of production $\phi_\beta$ and of degradation $\phi_\alpha$. The simulations are carried out using the following parameter values: $[XIST]_0 = 0$; $\Delta x = 0.1$; $\Delta t = 10^{-3}$; $n_\beta = 4$, $n_\alpha = 4$; $D_X = 1.0$; $\alpha_X = 1.0$; $\beta_X = 1.0$ (units arbitrary).

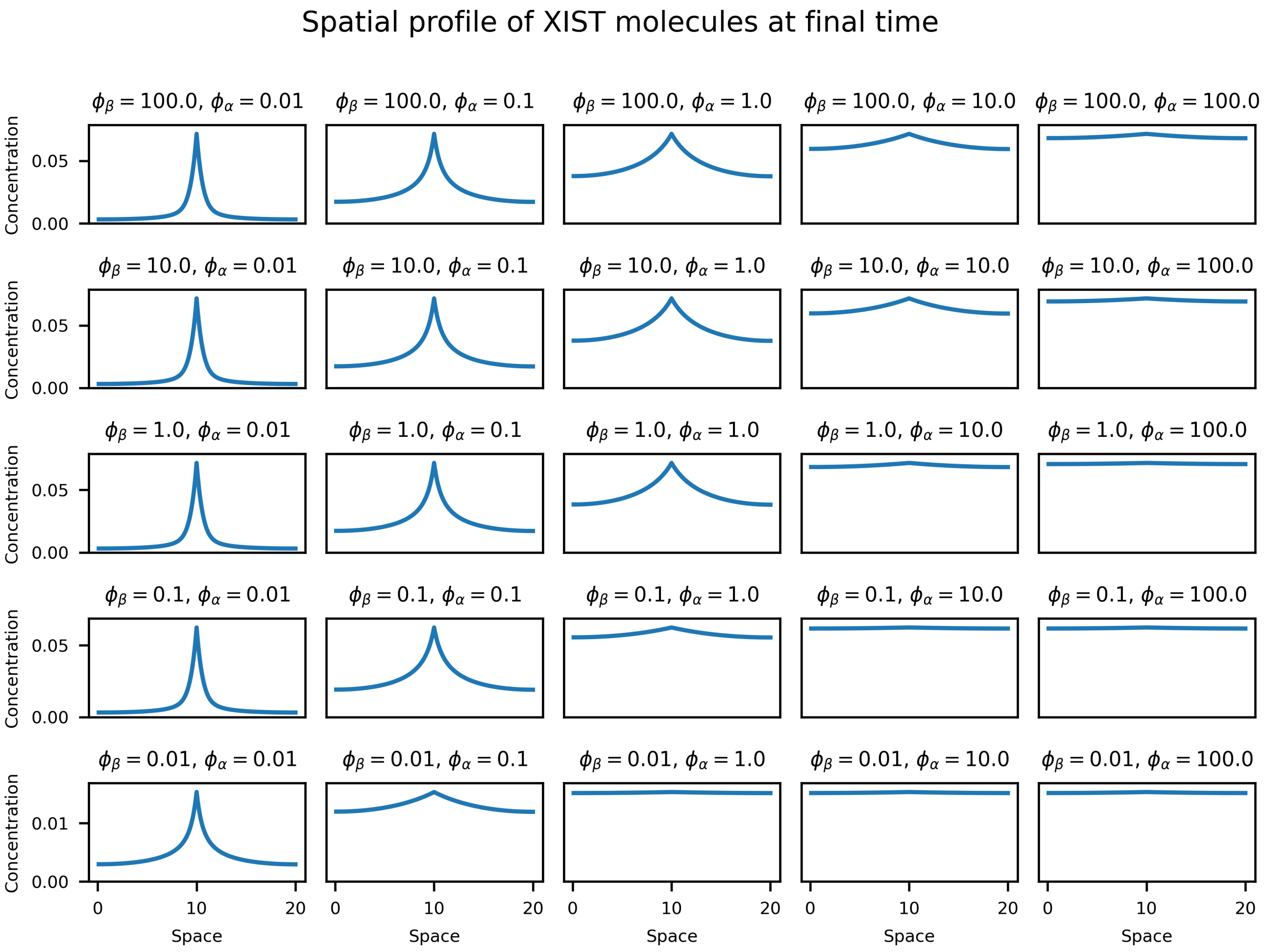

Fig. S2: Steady spatial profile of XIST concentration at t=2000 time units for different values of feedback coefficients of production $\phi_\beta$ and of degradation $\phi_\alpha$. The simulations are carried out using the parameter values used for Fig.S1

spreading takes place most rapidly and most homogeneously. To identify the optimal parameter set we have devised a scheme of bubble plot for the entire parameter scan. For each of the parameter set we have devised a bubble whose radius is proportional to the ratio of minima to the maxima of the XIST concentration throughout space at the asymptotic time limit. Moreover, the brightness of the color of the bubbles were determined by the speed of attainment of the steady value. More precisely, the brightness is directly proportional to the area under the curve depicting the variation of normalized spatially averaged XIST concentration with time. The entire scheme is illustrated in Fig.S3 for enhancing the clarity. This bubble visualization reveals that for $\phi_\alpha = 1.0$ and $\phi_\beta = 0.1$, the XIST molecules spread throughout the space with satisfactory temporal rapidness and spatial homogeneity.

##### 3.1.2 XCI (Chromatin) module

Subsequently, we studied the X chromosome inactivation by numerically simulating the PDEs described by Eqs.(S.2-S.3). Firstly, we performed the numerical continuation of the constituting ODE parts using $\delta$ as the free parameter of the PDEs to evaluate the steady states of the system. Then we curated appropriate steady states from the numerical continuation result. The result of the numerical continuation and curated steady states are delineated in Fig.S4. Thereafter, We used numerical techniques to simulate the PDEs associated to XCI module, similar to that of the XIST module. At the initial phase of the simulation, we have stimulated the system out of the steady state to initiate the inactivation process. The functional form of the stimulation can be written as follows:

$$f_s = \frac{s}{2}\sin^8\left(\frac{n_{\text{stim}}\pi x}{L}\right) \text{ for } (0 < t < t_1) \tag{S.8}$$

$$= \frac{s}{2}\cos\left(\frac{\pi(t-t_1)}{2(t_2-t_1)}\right)\sin^8\left(\frac{n_{\text{stim}}\pi x}{L}\right) \text{ for } (t_1 < t < t_2) \tag{S.9}$$

$$= 0 \text{ for } (t_2 < t) \tag{S.10}$$

We used this manual stimulation to conduct a phenomenological study of XCI in one spatial dimension. Later in this paper, the stimuli are governed by the spatiotemporal dynamics of the XIST molecules. Subsequently, using the aforementioned stimulation function, we simulated various scenarios of X chromosome inactivation, revealing numerous interesting features portrayed in Fig.S5. The parameter values used for the simulations are organized in the Table.S1.

Fig.S5(**A**) represent the unpinned wave. To further establish the robustness of the unpinning we widened the space keeping other parameters fixed. The unpinned waved thus obtained is illustrated

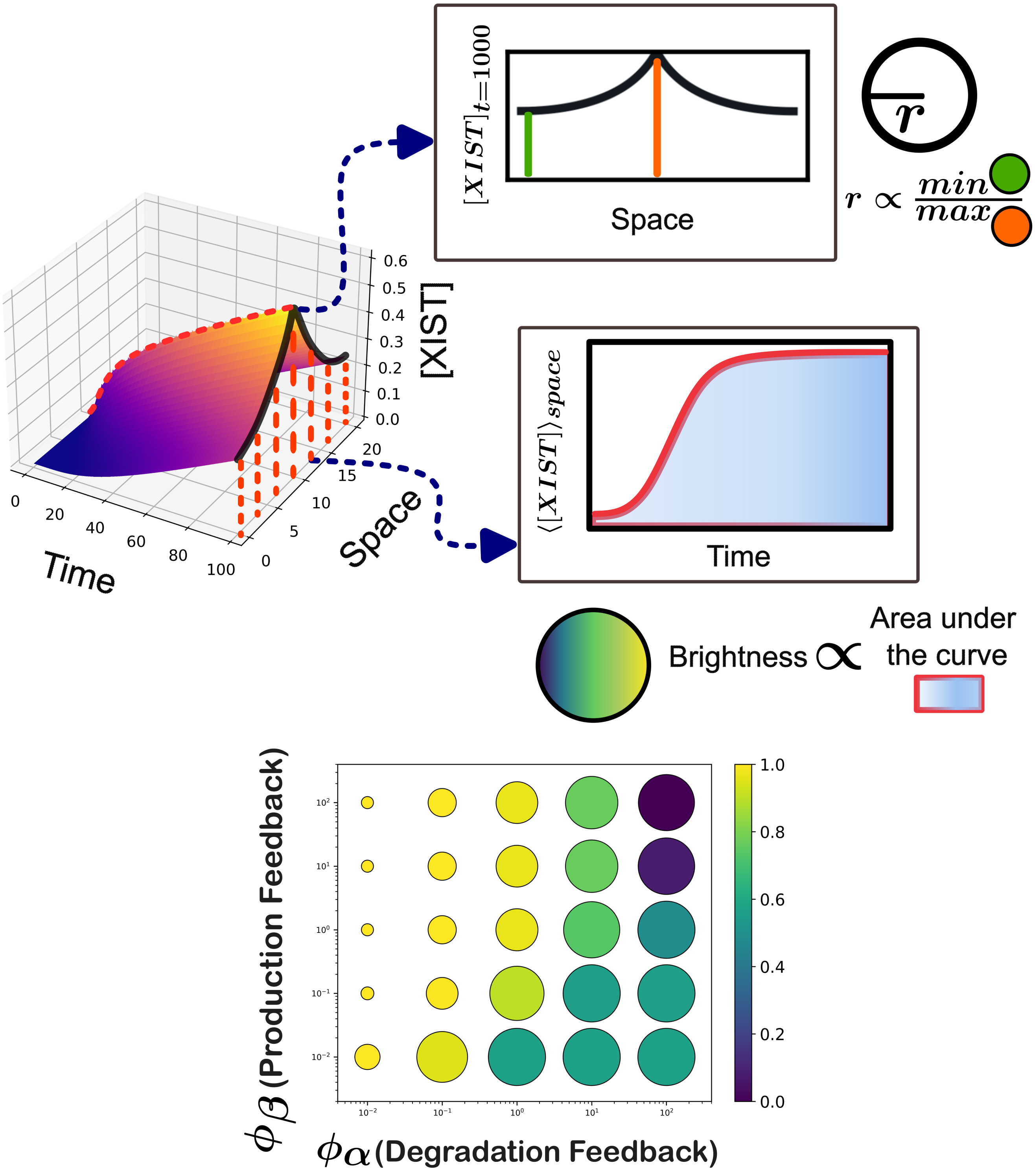

Fig. S3: The procedure for generating bubble plot from simulation data: Here each bubble represents the spatiotemporal profile of XIST concentration. The larger bubbles represent more spatial spreading of XIST molecules and brighter bubbles represents rapid attainment of steady spatial profile. The parameters are same as used in Fig.S1.

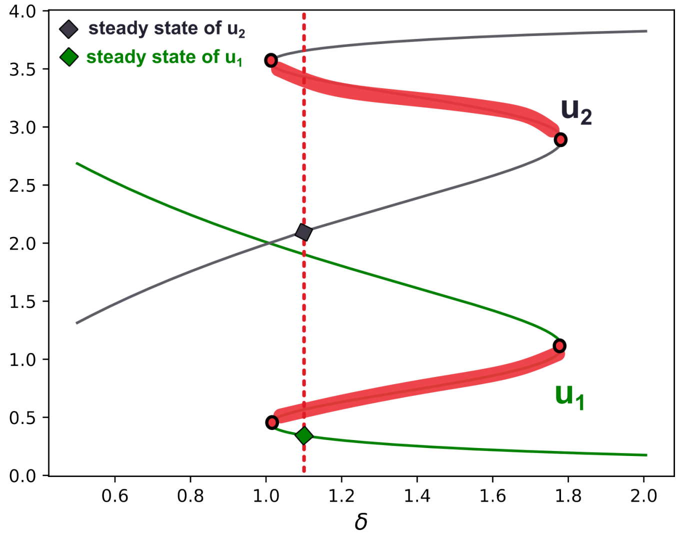

Fig. S4: Numerical continuation result of the XCI module. The parameters used are as follows: $\kappa_0 = 0.067$; $K = 1.0$; $\gamma = 1.0$; $n = 4$; $\alpha_{u_1} = \alpha_{u_2} = 0.05$; $\beta_{u_1} = \beta_{u_2} = 0.1$.

in Fig.S5(**B**) and the result implies that the unpinned state is truly unpinned arising out of the underlying dynamics of the system and it is independent of the length of simulated space.

Next we obtained the pinned wave from the unpinned wave in three ways and for each of them we tweaked only one of the parameters from the unpinned scenario. Fig.S5(**C**) represent the unpinned wave obtained by tweaking the feedback coefficient $\gamma$. Here we used $\gamma = 1$ for the region $2 < x < 18$ and $\gamma = 0.5$ otherwise. Thereafter, starting from the unpinned state of Fig.S5(**A**) we increased $\alpha_{u_1}$ and $\alpha_{u_2}$ from 0.05 to 0.06 keeping all other parameters fixed and that change in alpha value transformed the unpinned state into a pinned state and the pinned state is depicted in Fig.S5(**D**). Lastly, in Fig.S5(**E**) we illustrate the pinned state arising out of the increase in the inverse of feedback coefficient $K$ from 1.0 to 1.2.

Next, we studied the way to control the extent of inactivation and for that purpose we chose a relatively less parameter $K$ which modulates the feedback of the system. In Fig.S6 we portrayed the results for different $K$ values. The Figs.S6 (**A**), (**B**), and (**C**) portray the spatiotemporal profile of X inactivation respectively for $K = 1.0$, $K = 1.1$ and $K = 1.2$. The results unravel that with increase in $K$, i.e., the inverse of the positive feedback coefficient, the extent of X chromosome inactivation decreases.

##### 3.1.3 Effect of spatial constraint of chromatin on XCI

In the simulations carried out in previous subsections, we have assumed that the chromatin structure is continuously distributed throughout the one-dimensional space. Although this assumption

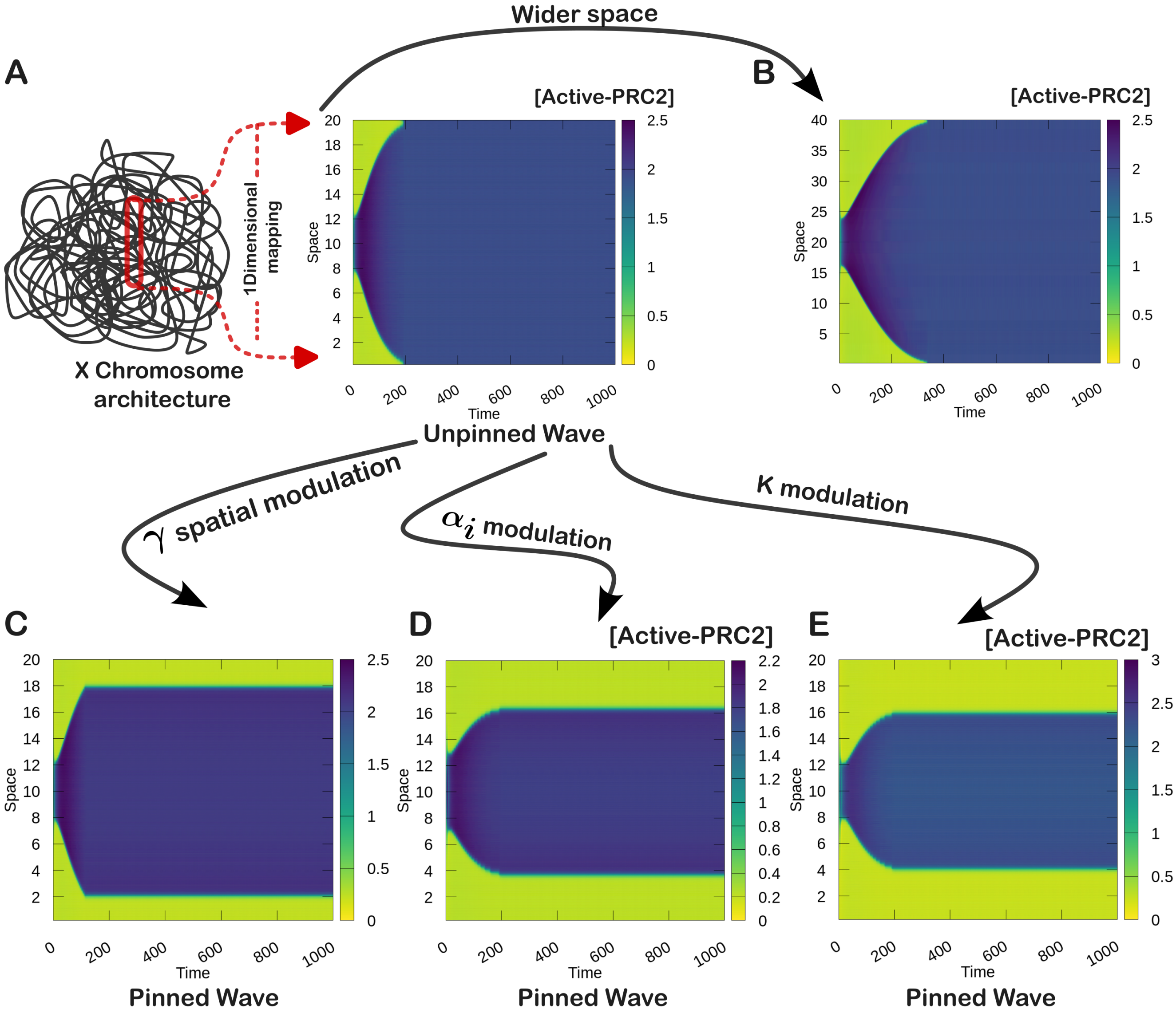

Fig. S5: Simulation result of XCI module for different parameter values: **(A)** Mapping of chromosomal structure in one dimension and the inactive wave propagation leading to unpinned wave for the parameters $u_1^0 = 0.3$; $u_2^0 = 2.0$; $\Delta x = 0.2$; $L = 20$; $\Delta t = 10^{-3}$; $D_{u_1} = 0.025$; $D_{u_2} = 10.0$; $\delta = 1.10$; $\kappa_0 = 0.067$; $K = 1.0$; $\gamma = 1.0$; $n = 4$; $\alpha_{u_1} = \alpha_{u_2} = 0.05$; $\beta_{u_1} = \beta_{u_2} = 0.1$; $n_{\text{stim}} = 1$; $t_1 = 2$; $t_2 = 4$. **(B)** The depiction of unpinned wave with parameters same as (A) but for a wider spatial region, i.e., $L = 40$. **(C)** Wave pinning achieved by spatial modulation of the parameter $\gamma$, other parameters are similar to that of (A). **(D)** Wave pinning attained by $\alpha_i$ modulation, here the parameters are same as (A) but for $\alpha_{u_1} = \alpha_{u_2} = 0.06$. (E) Wave pinning achieved by variation of $K$, parameters are same as (A) but for $K = 1.2$. (units arbitrary)

Table S1: Parameters used to obtain results portrayed in Fig.S5

| Figure | Parameters |
|---|---|
| Fig.S5 **(A)** | $u_1^0 = 0.3$; $u_2^0 = 2.0$; $\Delta x = 0.2$; $L = 20$; $\Delta t = 10^{-3}$;<br>$D_{u_1} = 0.025$; $D_{u_2} = 10.0$; $\delta = 1.10$; $\kappa_0 = 0.067$; $K = 1.0$; $\gamma = 1.0$; $n = 4$;<br>$\alpha_{u_1} = \alpha_{u_2} = 0.05$; $\beta_{u_1} = \beta_{u_2} = 0.1$; $n_{\text{stim}} = 1$; $t_1 = 2$; $t_2 = 4$ |
| Fig.S5 **(B)** | Same as **(A)** but for $L = 40$ |
| Fig.S5 **(C)** | Same as **(A)** but for $\gamma = 1$ at $(2 < x < 18)$ and $\gamma = 0$ otherwise |
| Fig.S5 **(D)** | Same as **(A)** but for $\alpha_{u_1} = \alpha_{u_2} = 0.06$ |
| Fig.S5 **(E)** | Same as **(A)** but for $K = 1.2$ |

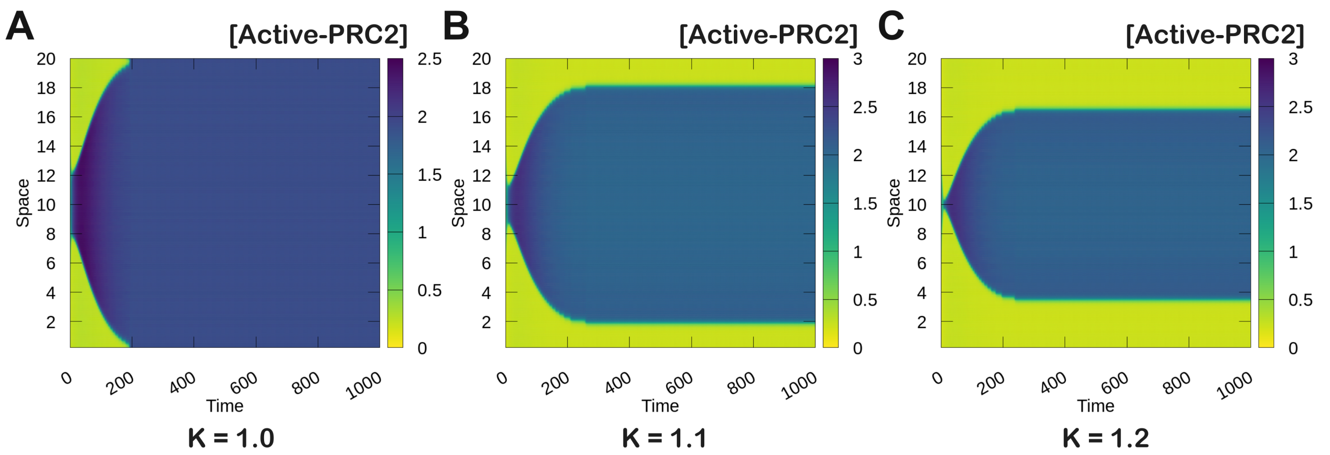

Fig. S6: Simulation result of XCI module for variation of the feedback coefficient $K$:**(A)** The parameter values are similar to Fig.S5**(A)**, i.e., $K = 1.0$; **(B)** same as **(A)** but for $K = 1.1$; **(C)** same as **(A)** but for $K = 1.2$. (units arbitrary)

is useful for exploring various dynamical features of the model, a more complete description requires considering the discrete nature of chromatin structure. In this subsection, we will examine the impact of the discrete nature of chromatin on the process of X chromosome inactivation.

In Figs.S8 and S9 we have portrayed different scenarios in terms of the discrete chromatin structures mapped in one spatial dimension. Here we have implemented the void space (white regions in our visualization) by distinctively assigning value of $\gamma$, i.e., $\gamma$ effectively can have two values 1 and 0 depending on the spatial coordinate. From biochemical perspective this means the elimination of chromatin originated positive feedback at void spaces. The detailed description of the parameter values used for simulation are given in Table.S2.

The results obtained for periodic and zero flux boundary condition reveals that the waves repeal each other and this repulsion is originated from the interplay between the active and inactive forms of PRC2. The detailed description of the spatiotemporal interaction that lead to the phenomenon of wave repulsion has already been discussed in the main text.

Moreover, the results also suggest that irrespective of the boundary conditions the steady spatial profile of the chromosomal activation is dependent on the final orientation of the chromatin structure. We have used this highly important feature of this model to justify our use of Xi chromosome structure instead of Xa structure for 3D simulations.

The compaction depicted in Figs.S8(**G**) and S9(**G**) respectively for periodic and zero-flux boundary conditions has been carried out by defining the void space at $t = 0$ by setting $\gamma = 1.0$ for $0, 30 < x < 10, 40$ and $\gamma = 0.0$ otherwise. During the numerical simulation, we implemented the compaction by redefining the void space at each $1200^{th}$ time steps, i.e., at the $i * 1200th$ time step ($i =$ integer) of the simulation the void space is defined as $\gamma = 1.0$ for $0+i, 30-i < x < 10+i, 40-i$ and $\gamma = 0.0$ otherwise. The compaction stops when the terminal regions of the chromatin reaches a predefined point in space. Furthermore, to ensure the conservation of mass during this whole compaction process, we have mapped the concentration of $u_1$ and $u_2$ at each compaction step which we have termed as state mapping. The detailed procedure for state mapping during compaction is illustrated in Fig.S7.

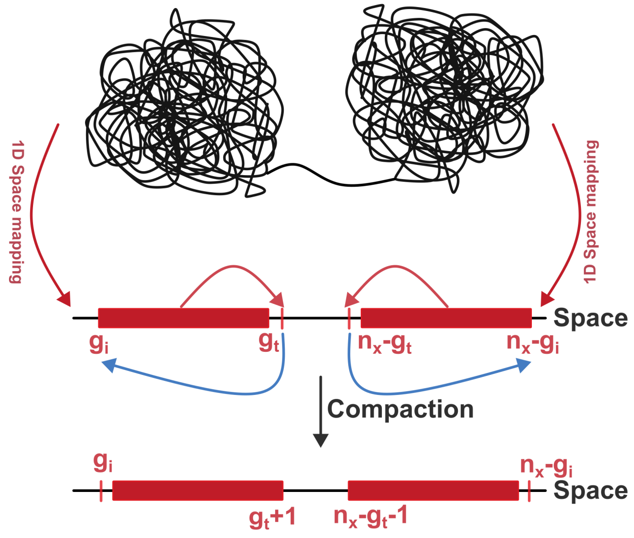

Fig. S7: The schematic diagram depicting details of state mapping during compaction.

The state mapping has been implemented as follows:

$$\begin{aligned}
&\text{mem} = u_j(g_t+1);\ u_j(g_i+1:g_t+1) = u_j(g_i:g_t); u_j(g_i) = \text{mem} \\
&\text{mem} = u_j(n_x-(g_t+1)); \\
&u_j(n_x-(g_t+1):n_x-(g_i+1)) = u_j(n_x-g_t:n_x-g_i);\ u_j(n_x-g_i) = \text{mem}
\end{aligned} \tag{S.11}$$

Where, $g_i$ and $g_t$ are indices defining the edges of the chromatin region at $t = 0$ and $n_x$ is the total number of spatial grid points. mem is a dummy variable used during the process.

###### 3.1.4 Integrating XIST module with XCI module through Tethering module

Subsequently, we integrated the XIST module and the X chromosome inactivation module by the tethering module. The governing principles of the tethering module have been provided in Sec.2.3. The simulation results are depicted and discussed in the main text. We have provided an elaborated account of the parameters used to get the results in Table.S3.

##### 3.2 Simulations performed in three dimensions

In the next stage of our study we explored the whole phenomenon of X chromosome inactivation using a reasonable three dimensional X chromosome architecture. Using the continuum model was the most significant bottleneck for simulating the model in three dimensions with discrete

Table S2: Parameters used to obtain results portrayed in Fig.S8 and Fig.S9

| Figure | Parameters (units arbitrary) |
|---|---|
| Fig.S8 **(A)** | $u_1^0 = 0.2683$; $u_2^0 = 2.0$; $\Delta x = 0.2$; $L = 40$; $\Delta t = 10^{-3}$;<br>$D_{u_1} = 0.025$; $D_{u_2} = 10.0$; $\delta = 1.20$; $\kappa_0 = 0.067$; $K = 1.1$; $n = 4$;<br>$\alpha_{u_1} = \alpha_{u_2} = 0.05$; $\beta_{u_1} = \beta_{u_2} = 0.1$; $n_{\text{stim}} = 1$; $t_1 = 2$; $t_2 = 4$<br>$\gamma = 1.0$ (for, $0 < x < 10$) and $= 0.0$ otherwise; Boundary= Periodic |
| Fig.S8 **(B)** | Same as Fig.S8**(A)** but for:<br>$\gamma = 1.0$ (for, $5 < x < 15$) and $= 0.0$ otherwise |
| Fig.S8 **(C)** | Same as Fig.S8**(A)** but for:<br>$\gamma = 1.0$ (for, $10 < x < 20$) and $= 0.0$ otherwise |
| Fig.S8 **(D)** | Same as Fig.S8**(A)** but for:<br>$\gamma = 1.0$ (for, $0 < x < 10$, $30 < x < 40$) and $= 0.0$ otherwise |
| Fig.S8 **(E)** | Same as Fig.S8**(A)** but for:<br>$\gamma = 1.0$ (for, $5 < x < 15$, $25 < x < 35$) and $= 0.0$ otherwise |
| Fig.S8 **(F)** | Same as Fig.S8**(A)** but for:<br>$\gamma = 1.0$ (for, $9 < x < 19$, $21 < x < 31$) and $= 0.0$ otherwise |
| Fig.S8 **(G)** | Same as Fig.S8**(A)** but for:<br>$\gamma = 1.0$ (for, $0 < x < 10$, $30 < x < 40$) and $= 0.0$ otherwise initially,<br>The chromatin regions move one grid every 1200th time step till:<br>$\gamma = 1.0$ (for, $9 < x < 19$, $21 < x < 31$) and $= 0.0$ otherwise |
| Fig.S9 **(A)**-**(G)** | Same as Fig.S8 **(A)**-**(G)** but for:<br>Boundary= Zero-Flux |

Table S3: Parameters used to obtain results of integrated module (Figure 4)

| Modules | Parameters (units arbitrary) |
|---|---|
| **Simulation** | $\Delta x = 0.2$; $\Delta t = 10^{-3}$; $L = 80$; Boundary= Zero-Flux |
| **XIST** | $[XIST]_0 = 0$; $n_\beta = 4$; $n_\alpha = 4$; $D_X = 1.0$; $\alpha_X = 0.01$;<br>$\beta_X = 1.0$ (for $x = 40$) and $= 0.0$ (otherwise);<br>$\phi_\beta = 0.1$; $\phi_\alpha = 1.0$; |
| **XCI** | $u_1^0 = 0.3$; $u_2^0 = 2.0$; $D_{u_1} = 0.025$; $D_{u_2} = 10.0$; $\delta = 1.10$;<br>$\kappa_0 = 0.067$; $K = 1.2$; $\gamma = 1.0$; $n = 4$; $\alpha_{u_1} = \alpha_{u_2} = 0.05$;<br>$\beta_{u_1} = \beta_{u_2} = 0.1$ |
| **Tethering** | $n_{\text{thr}} = 10$; $n_{\text{total}} = 400$; $\phi_{\text{thr}} = 0.5$ |

### Periodic Boundary Condition

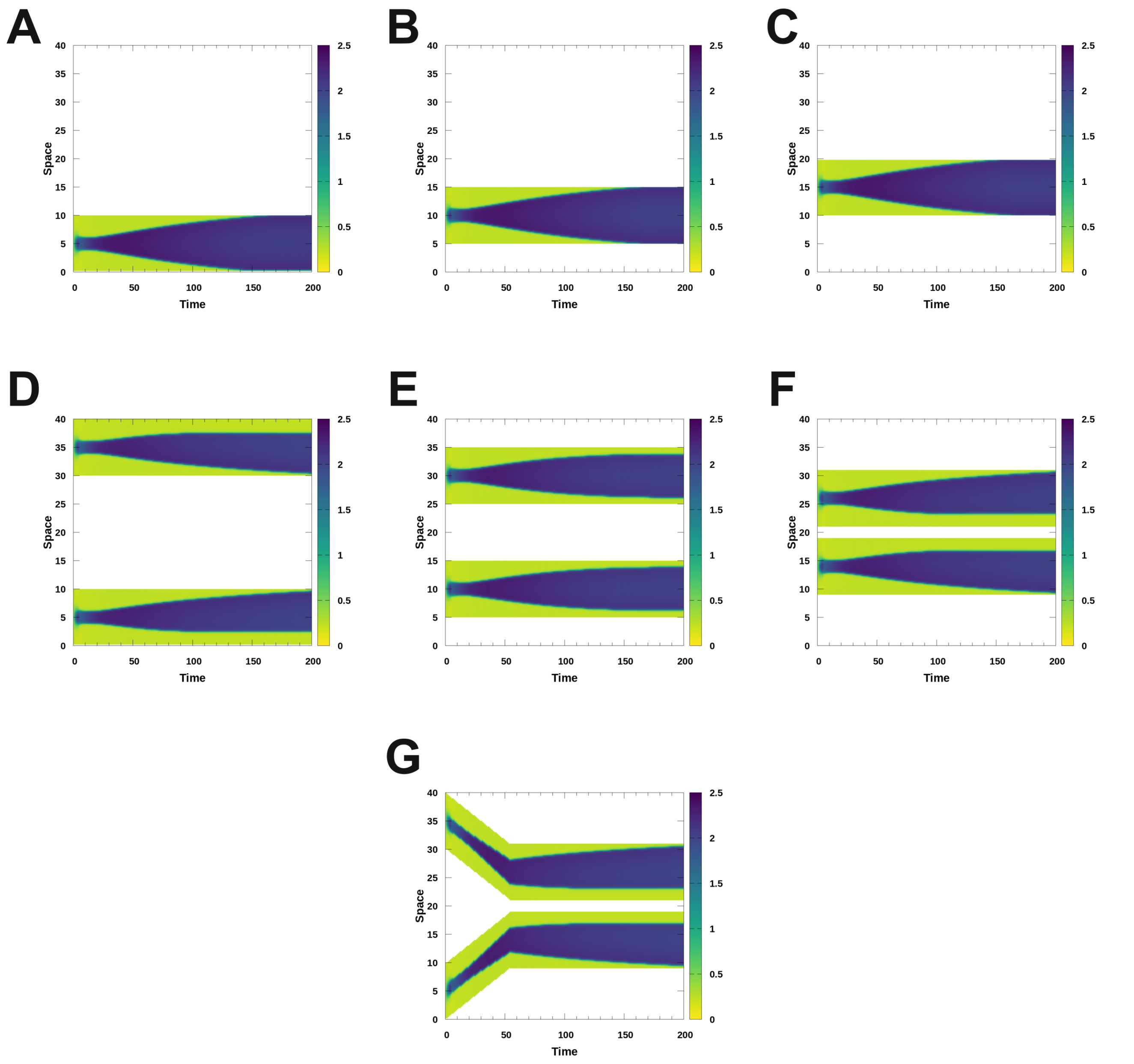

Fig. S8: Effect of spatial constraint on the process of XCI for **periodic boundary condition**. An itemized description of the parameters used and other specification of the simulations are provided in Table.S2.

### Zero-Flux Boundary Condition

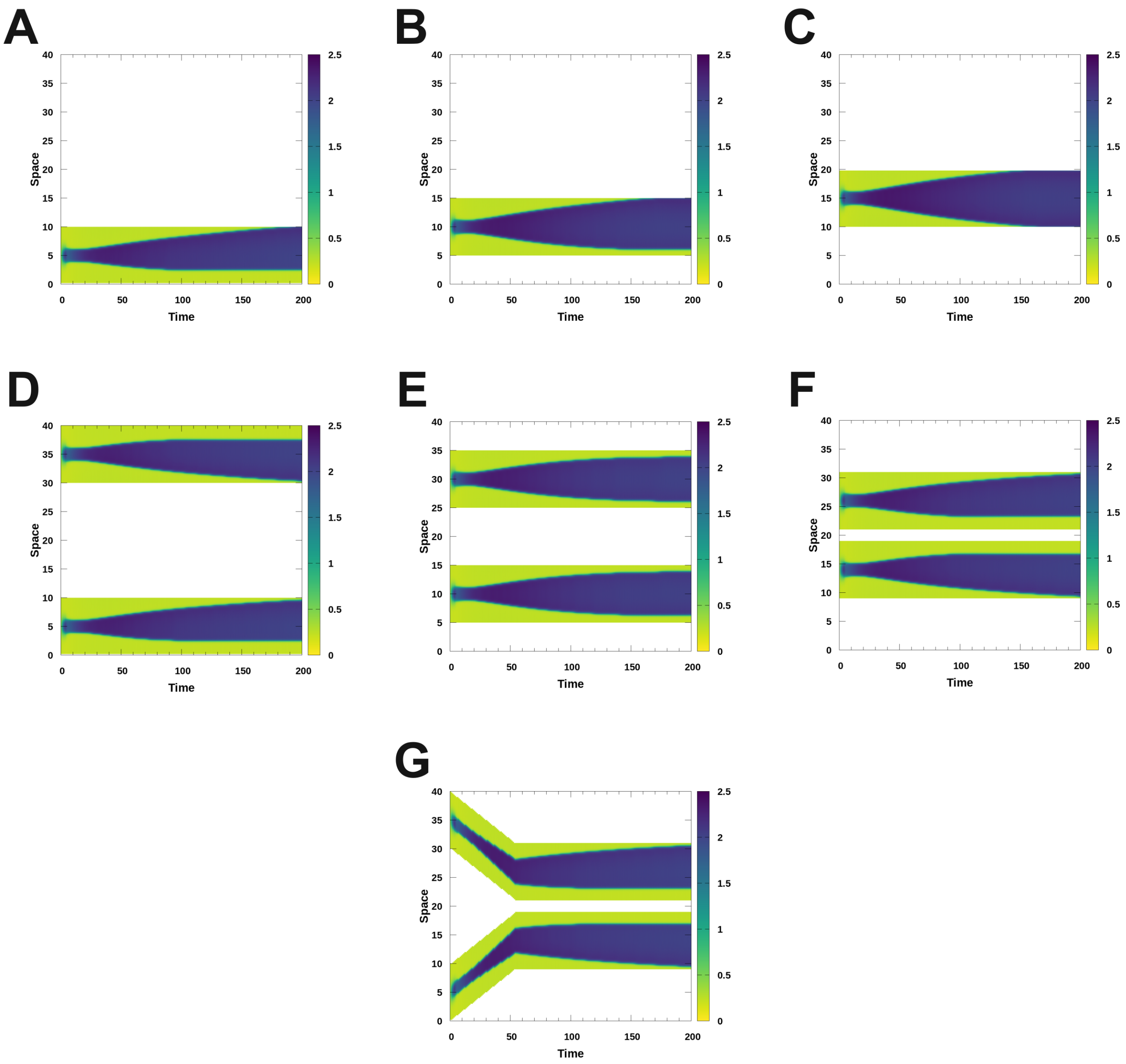

Fig. S9: Effect of spatial constraint on the process of XCI for **zero-flux boundary condition**. An itemized description of the parameters used and other specification of the simulations are provided in Table.S2.

chromosome structure. The main challenge was to include the effect of chromosome structure into the spatiotemporal dynamics of X chromosome inactivation. We have overcome this obstacle by spatially modulating the feedback parameter $\gamma$, which is highly sensitive parameter towards the bi-stability of the system. As the bi-stability is the key tenet of the system to exhibit wave pinning, with diminished $\gamma$ the system effectively gets converted to a mono-stable one. For the three dimensional model we have made $\gamma$ space dependent, i.e., for each spatial point we calculated the lowest distance of that particular point from the chromosome architecture and then defined the effective $\gamma$ for that point as follows:

$$\gamma(x, y, z) = \gamma_0 e^{-\kappa d} \tag{S.12}$$

This suggests that both the feedback and the bi-stability of the system are influenced by the chromosome structure. For $d = 0$, where the spatial points reside on the chromosome structure, $\gamma = \gamma_0$. For very large $d$ values, where the spatial points are far from the chromosome structure, $\gamma \approx 0$. This dependence of $\gamma$ on the chromosome structure incorporates the essential influence of the chromosome's structure into the system's spatiotemporal landscape. Additionally, this approach to $\gamma$ captures the transient movements of the chromosome in space and considers the structure as an averaged one. This is justified, as we have used the semi-empirical chromosome architecture derived from HiC data [8] and interpolated the chromosome structure to improve gene resolution. Moreover, the $\gamma$ spatial modulation ensures that the $\gamma$ on an average takes higher value for chromatin dense regions leading to bi-stability and chromosomal inactivation and vice versa. To facilitate reader convenience and clarity, we present the necessary details for the 3D simulation in an itemized format in the following text box.

**3D simulation details**

The entire procedure of 3D simulation is provided in a chronological order as follows:

1. We obtain the Cartesian co-ordinates (700 data points) of the chromosome structure from [8] and then we interpolate those 700 points to generate 20000 overall data points. In reality, the X chromosome has approximately 155 million base pairs and the increase in structural resolution of X chromosome later facilitated the comparison of simulated result with reported experimental data by enhancing the gene resolution.

2. We then discretized the space according to the coordinates of the X chromosome structure and that resulted in a simulation volume in the shape of a parallelepiped. The details of the dimension in provided later in the Table.S4 containing the parameter

details.

3. Subsequently, we define the space dependent $\gamma$ values according to the Eq.S.12 for XCI module and for the XIST module we fixed the X inactivation center , i.e., the point in 3D space with nonzero $\beta_X$ value.

4. Thereafter, we utilized explicit Euler algorithm to simulate the PDEs described by Eqs.S.1, S.2 and S.3. We have implemented the tethering of XIST molecules following the conditions described in Eq.S.5. The detailed description of various parameters used are presented in Table.S4.

5. Next, we map the x-linked genes of a particular species (mouse, in the context of this paper) to the previously mentioned polymeric structure with 20000 segments and mark them as on or off according to the experimental results [9]. Lets take a test case to explain the procedure: The length of the mouse genome is ≈169 Mbp [10] which implies that each segment of the used polymeric structure correspond to ≈8473 bp. The XIST gene for mouse is located at the region 102503979 to 102526839 [10] and mapping of the XIST gene to the polymeric structure reveals that the XIST gene is located in between 12097 th and 12099 th segment of the same. Similarly, we have mapped all the genes that are known to escape the process of inactivation and tried to capture as many of them as possible through the simulation.

6. During the simulation we have not allowed the tethering of XIST molecules inside the sphere of radius 30 spatial units, centered around the XIC (12098 th segment of the polymeric chain).

7. Finally, we analyzed the simulation results and compared them with the experimental observations.

Table S4: Parameters used to obtain results of 3D module (Figure 5)

| Modules | Parameters (units arbitrary) |
|---|---|
| **Simulation** | $\Delta x = \Delta y = \Delta z = 0.5$; $\Delta t = 10^{-3}$; $L_x = 70$; $L_y = 72.5$; $L_z = 70.5$; Boundary= Zero-Flux |
| **XIST** | $[XIST]_0 = 0$; $n_\beta = 4$; $n_\alpha = 4$; $D_X = 25.0$; $\alpha_X = 0.01$; $\beta_X = 10.0$ (for XIC) and $= 0.0$ (otherwise); $\phi_\beta = 0.01$; $\phi_\alpha = 1.0$; |
| **XCI** | $u_1^0 = 0.3$; $u_2^0 = 2.0$; $D_{u_1} = 0.05$; $D_{u_2} = 10.0$; $\delta = 1.15$; $\kappa_0 = 0.067$; $K = 1.0$; $\gamma_0 = 1.0$; $\kappa = 0.5$; $n = 4$; $\alpha_{u_1} = \alpha_{u_2} = 0.05$; $\beta_{u_1} = \beta_{u_2} = 0.1$ |
| **Tethering** | $n_{\text{thr}} = 100$; $n_{\text{total}} = 20000$; $\phi_{\text{thr}} = 0.005$ |

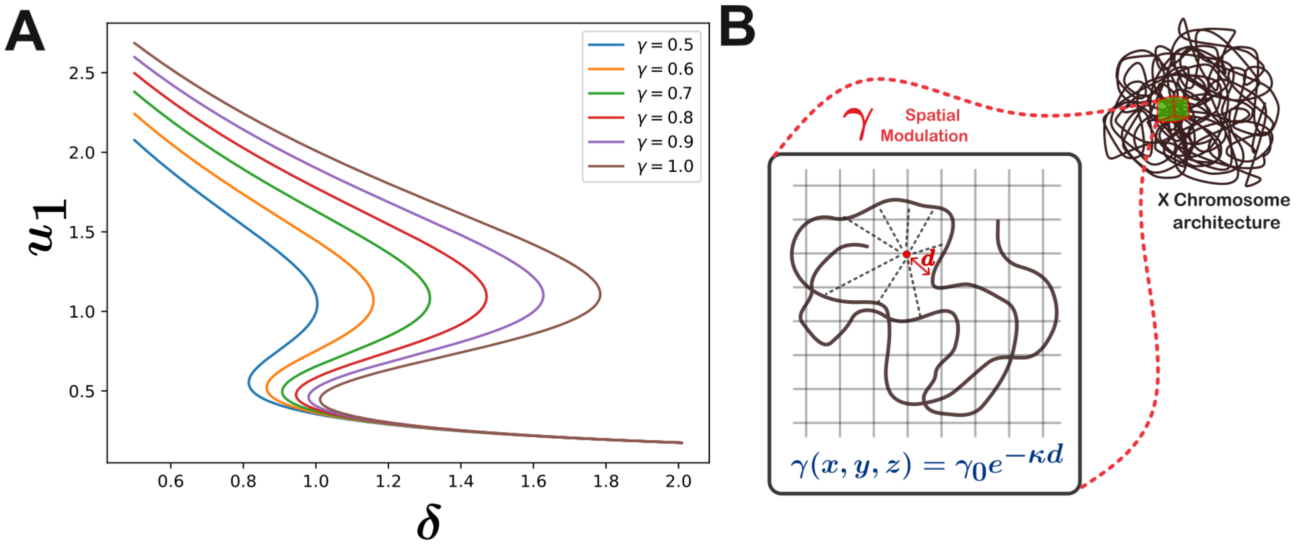

Fig. S10: **(A)** Depiction of variability of the bi-stability with variation in $\gamma$. The result implies high sensitivity of the parameter $\gamma$ on bi-stability. **(B)** A schematic diagram roughly illustrating the method of the spatial modulation of $\gamma$.

#### Insights on length and time scale of the 3D simulation

The typical length scale of the X chromosome in mammalian cells is approximately 1 $\mu$m [11]. In our 3D simulation framework, the average length of the simulated box is approximately 70 spatial units, meaning one spatial unit corresponds to 0.014 $\mu$m of physical length. Since XIST is the key driver of X chromosome inactivation dynamics, we normalized the time with one thousandth of the median half-life of XIST RNA molecules, which is approximately 0.004 hours or 14.4 seconds [12]. This normalization implies that one unit of the diffusion coefficient in our simulation corresponds to approximately $1.36 \times 10^{-5} \mu m^2/sec$ in physical units. For numerical reasons, we used a diffusion coefficient of XIST molecules $D_X = 25$, which corresponds to approximately $3.4 \times 10^{-4} \mu m^2/sec$. The sub-nuclear diffusion coefficient of mRNA molecules ranges from $5 \times 10^{-3}$ to greater than 1 $\mu m^2/sec$ [13]. Considering that lncRNA are much heavier than mRNA molecules, the diffusion coefficient of XIST used in our simulation falls within a reasonable range, being less than the lower bound of the diffusion coefficient of mRNA molecules.

#### 3.3 Analysis of 3D simulation data and comparison with experimental results

Once we obtained the simulation the new reasonable step was to analyze the data and compare the result with experimental results. We have used the gene expression data reported in Ref.[9] and mapped the genes in our chromosomal structure. We have considered a gene as escapee if it's expression probability is greater than 0.1. The result thus obtained is portrayed in Figure 5. Once we have the experimental results mapped to the chromosomal structure that we have used, we compare the simulation results with experimental results. We have categorized a gene as active according to the simulation, if there is at least 50 percent agreement between the simulation result and the experimental result. A detailed illustration of the condition is provided in Fig.S11. The analysis of the results are provided in the main text.

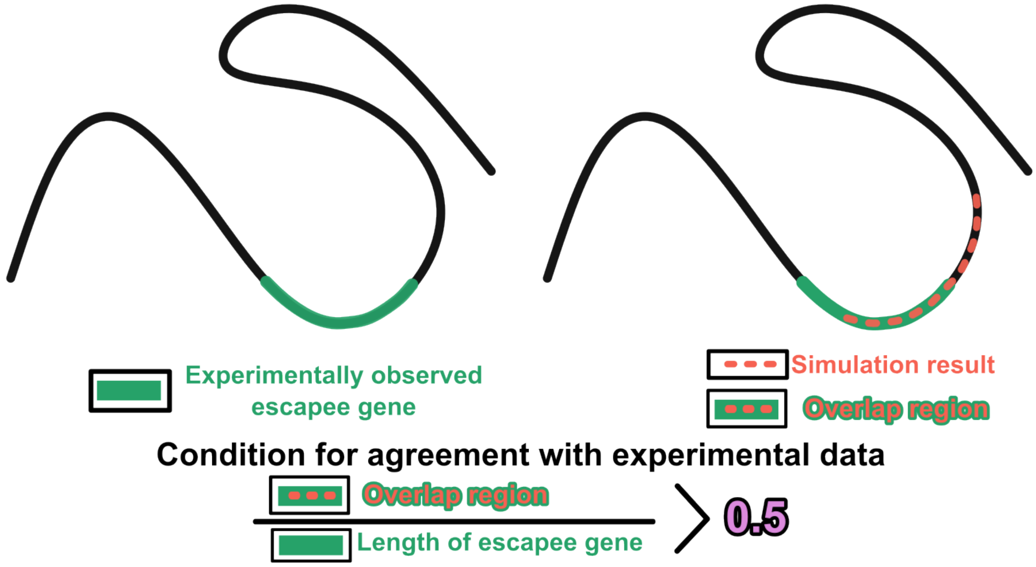

Fig. S11: Schematic representation of the condition for agreement between simulated and experimental results.
